# A brown fat-enriched adipokine, ASRA, is a leptin receptor antagonist that stimulates appetite

**DOI:** 10.1101/2023.09.12.557454

**Authors:** Lei Huang, Pengpeng Liu, Yong Du, J. Fernando Bazan, Dongning Pan, Qingbo Chen, Alexandra Lee, Vijaya Sudhakara Rao Kola, Scot A. Wolfe, Yong-Xu Wang

## Abstract

The endocrine control of food intake remains incompletely understood, and whether the leptin receptor (LepR)-mediated anorexigenic pathway in the hypothalamus is negatively regulated by a humoral factor is unknown. Here, we identify an appetite-stimulating factor – ASRA – that represents a peripheral signal of energy deficit and orthosterically antagonizes LepR signaling. *Asra* encodes an 8 kD protein that is abundantly and selectively expressed in adipose tissue and to a lesser extent, in liver. ASRA associates with autophagy vesicles and its secretion is enhanced by energy deficiency. In vivo, fasting and cold stimulate *Asra* expression and increase its protein concentration in cerebrospinal fluid. *Asra* overexpression attenuates LepR signaling, leading to elevated blood glucose and development of severe hyperphagic obesity. Conversely, either adipose- or liver-specific *Asra* knockout mice display increased leptin sensitivity, improved glucose homeostasis, reduced food intake, resistance to high-fat diet-induced obesity, and blunted cold-evoked feeding response. Mechanistically, ASRA acts as a high affinity antagonist of LepR. AlphaFold2-multimer prediction and mutational studies suggest that a core segment of ASRA binds to the immunoglobin-like domain of LepR, similar to the ‘site 3’ recognition of the A-B loop of leptin. While administration of recombinant wild-type ASRA protein promotes food intake and increases blood glucose in a LepR signaling-dependent manner, point mutation within ASRA that disrupts LepR-binding results in a loss of these effects. Our studies reveal a previously unknown endocrine mechanism in appetite regulation and have important implications for our understanding of leptin resistance.

---

Energy homeostasis is largely controlled by communications between the peripheral tissues and the central nervous system. In response to changes in the body’s energy reserves and nutritional status, peptides are secreted from peripheral tissues to regulate signaling pathways in the hypothalamus and brainstem, which in turn modulate food intake and body weight. Indeed, a number of peripheral anorexigenic peptides have been identified^1–5^. In contrast, very few peripheral orexigenic peptides have been identified^6, 7^ and how peripheral tissues communicate with central regulatory system to stimulate appetite was less understood.

The LepR signaling pathway is considered the most critical anorexigenic pathway for controlling food intake, energy balance, and body weight^8–13^. Leptin and its receptor (LepRb) exert the effects on food intake and body weight mainly through activation of the JAK2-STAT3 axis in pro-opiomelanocortin (POMC)-expressing neurons and Agouti-related protein (AgRP)-expressing neurons within the arcuate nucleus (ARC) of the hypothalamus. LepR signaling also regulates glucose metabolism in POMC neurons independently of food intake and body weight^8–13^. While leptin is an effective treatment for complete or partial leptin deficiency, common forms of obesity, including diet-induced obesity, are associated with elevated endogenous leptin levels, and show an inadequate response to exogenous leptin treatment, a state characterized as leptin resistance^8–12^. Elucidating the underlying mechanisms of leptin resistance may identify regulatory pathways that can be used to modify common metabolic disorders. To date, whether peripheral tissues produce endocrine factors to attenuate LepR signaling that may exacerbate leptin resistance remains undefined.

## Results

### Identification of a brown fat (BAT)-enriched adipokine that is induced by fasting and cold

Given that leptin is selectively expressed in white fat (WAT) and that cold^14–17^ and robust WAT browning^18^ induce hyperphagia, we considered the possibility that a BAT-enriched adipokine might exist to counterbalance leptin signaling. We further envisioned that such an adipokine is likely upregulated during fasting. We therefore analyzed gene expression datasets^19, 20^ and screened for genes that are abundantly and selectively expressed in BAT and are induced by fasting (**Extended Data Fig. 1a**). Bioinformatic tools^21, 22^ were then used to predict secreted proteins. These combined analyses identified a previously uncharacterized mouse gene *1190005I06RIK*, and its human ortholog *C16orf74*. *1190005I06RIK* gene has an open reading frame (ORF) of 111 amino acids, while *C16orf74* has an ORF of 76 amino acids (**Extended Data Fig. 1b**). However, the *1190005I06RIK* mRNA has an internal in-frame ATG that corresponds to the start codon of *C16orf74* and this internal ATG is flanked by a strong Kozak sequence (GACGCC<u>ATG</u>G), raising the possibility that this represents a key translation initiation site. Indeed, two bands, 12 kD and 8 kD, were detected in HEK293 cells transfected with expression plasmids containing the mouse 111-amino acid ORF. The 8 kD band was the product of translation initiated from the internal ATG, as it disappeared when the corresponding Methionine residue was mutated to Alanine (**Extended Data Fig. 1c**). Importantly, only the 8 kD product was detected in mouse adipose tissue and adipocyte culture (**Extended Data Fig. 1d and 1e**). Thus, translation of endogenous *1190005I06RIK* mRNA is initiated from the internal in-frame ATG, producing an 8 kD polypeptide with 76 amino acid residues highly homologous to human *C16orf74*. Neither of these proteins possesses a signal peptide, but both are predicted by SecretomeP^22^ to be non-classically secreted proteins with high NN-scores that are similar to that of fibroblast growth factor 1 (FGF1), a non-classically secreted protein. Based on its tissue expression and functional analysis described below, we named this gene as *Asra* (<u>a</u>dipose-<u>s</u>ecreted regulator of <u>a</u>ppetite). cDNA constructs encoding the 76-amino acid polypeptide were used in our studies.

*Asra* was selectively expressed in adipose tissue and to a lesser extent, in the liver (**Fig. 1a**). In adipose tissue, *Asra* was expressed in mature adipocytes (**Extended Data Fig. 1e and 1f**). We confirmed that expression of *Asra* in both adipose and liver was induced by fasting and repressed by re-feeding (**Fig. 1b**). Expression of *Asra* was also upregulated by cold in these tissues (**Fig. 1c and Extended Data Fig. 1g**). Thus, *Asra* expression is induced by conditions that promote a negative energy balance. In addition, liver *Asra* expression was associated with high-fat diet feeding (**Fig. 1d**), while adipose *Asra* expression was not significantly altered (**Extended Data Fig. 1h**).

**Figure 1.**
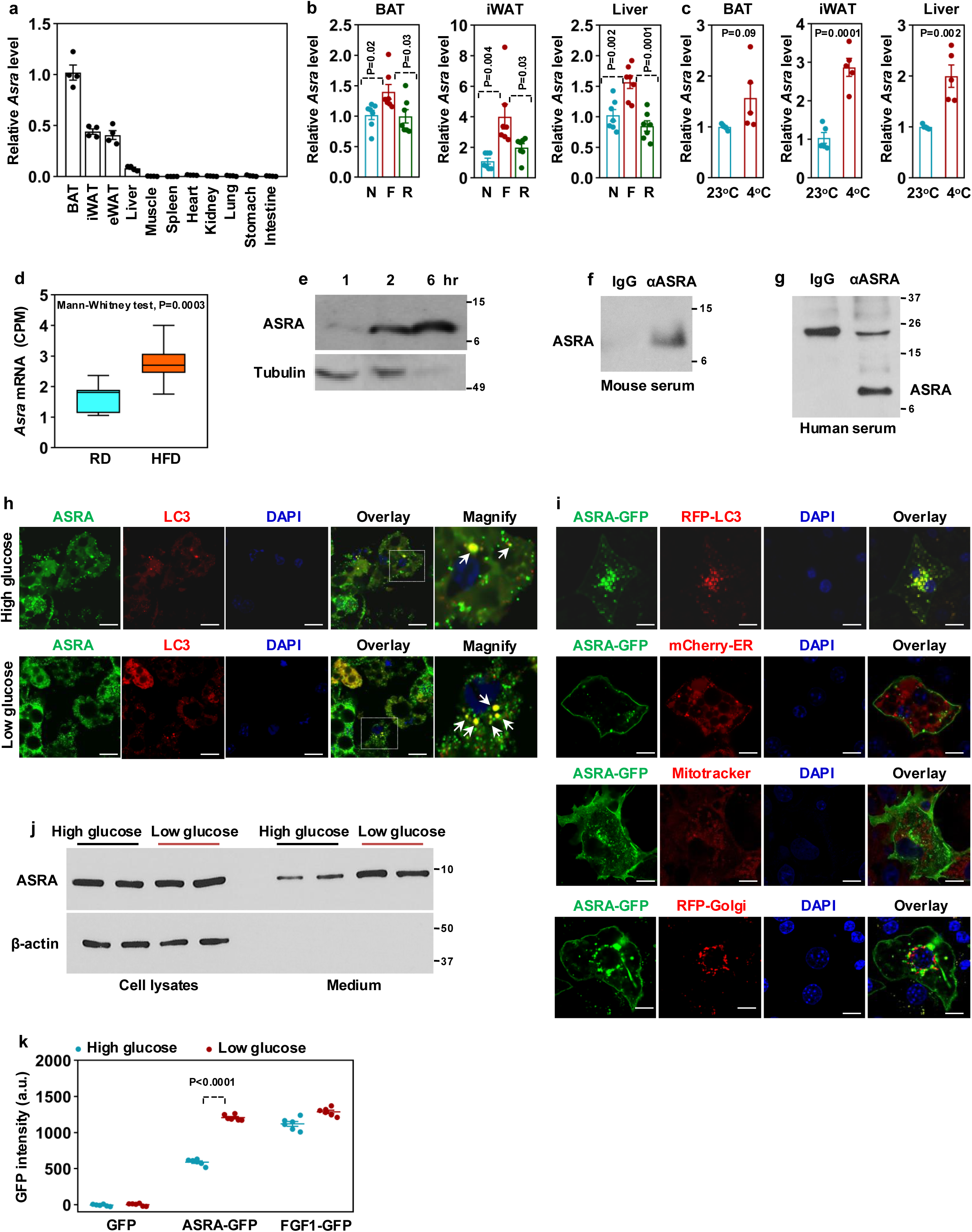
Identification of ASRA as a BAT-enriched adipokine induced by fasting and cold and associated with autophagy vesicles. **a,** *Asra* mRNA expression in mouse tissues (n=4). **b,** *Asra* mRNA expression in BAT, iWAT, and liver of mice subjected to non-fasting (N), 12-hr fasting (F), and 3-hr re-feeding (R). n=7/group. **c,** *Asra* mRNA expression in BAT, iWAT, and liver of mice subjected to 8-hr cold exposure. n=5/group. **d,** *Asra* mRNA expression in liver of mice fed with a regular diet (RD) (n=11) or a high-fat diet (HFD) for three weeks (n=10). The RNA-seq data were downloaded from GSE88818 dataset. **e,** ASRA secretion from BAT ex vivo. **f, g,** Detection of ASRA in mouse serum (**f**) or human serum (**g**) after immunoprecipitation with ASRA antibodies. **h,** Co-localization of endogenous ASRA and LC3 in mature adipocytes cultured in FBS-free DMEM medium containing high glucose (4.5 g/L) or low glucose (1 g/L) for 6 hr. Bar=50 μm. **i,** ASRA-GFP localization in mature adipocytes. Bar=50 μm. **j,** ASRA secretion from mature adipocytes cultured in FBS-free DMEM medium containing high glucose or low glucose for 6 hr. **k,** ASRA-GFP secretion from HEK293 cells cultured in FBS-free DMEM medium containing high glucose or low glucose for 6 hr (n=6).

Analysis of microarray data^23^ of subcutaneous fat found that levels of ASRA were positively correlated with body mass index (BMI) in a large cohort of 770 men (**Extended Data Fig. 1i**). In addition, genome-wide association studies have found genetic variants of ASRA that loosely associated with type 2 diabetes in both Japanese population (rs3777457, p=6 × 10^−7^)^24^ and European population (rs439967, p=2 × 10^−6^)^25^.

Cultured adipocytes secreted ASRA into conditioned medium (**Extended Data Fig. 1j**). To examine ASRA secretion ex vivo, small pieces of adipose tissue were incubated with conditioned medium. While tubulin was detected in the medium in the first two hours due to initial tissue breakage, ASRA was more abundantly present at the 6-hr time point (**Fig. 1e**), indicating an active secretion. ASRA was also detected in mouse and human serum after enrichment with immunoprecipitation (**Fig. 1f and 1g**).

### ASRA associates with autophagy vesicles and its secretion is induced by starvation

To gain information of how ASRA is secreted, we examined its subcellular localization. Immunofluorescent staining of endogenous ASRA in adipocytes revealed that ASRA was localized in vesicles and cell periphery (**Fig. 1h**). On the basis that secretory autophagy pathway is one route for non-classical protein secretion^26–28^ and that *Asra* expression was induced by fasting, we examined whether ASRA associates with autophagy vesicles. We found that some ASRA-containing vehicles were also positive for the autophagy marker LC3, and the number of these double-labeled vesicles was increased under conditions of low glucose (**Fig. 1h**), indicating that ASRA vesicles evolve into or fuse with autophagy vesicles during energy deficiency. To further validate this observation, we expressed a fusion of GFP with the C-terminus of ASRA in adipocytes. Similar to endogenous ASRA, ASRA-GFP displayed a vesicular and cell peripheral localization pattern (**Fig. 1i**). The ASRA-GFP marked vesicles were distinct from the membrane structures of endoplasmic reticulum, Golgi, and mitochondria. Instead, they almost completely overlapped with RFP-LC3 marked-vesicles (**Fig. 1i**). In contrast, FGF1-GFP was present in the cytoplasm and cell periphery as reported^29^ (**Extended Data Fig. 1k**). Association of ASRA with autophagy vesicles was not cell type-specific, as similar results were also observed in COS7 cells (**Extended Data Fig. 1l**). Interestingly, low glucose medium enhanced the secretion of ASRA from adipocytes (**Fig. 1j**). We also compared ASRA-GFP and FGF1-GFP secretion in HEK293 cells by measuring GFP fluorescence intensity in medium. Secretion of ASRA-GFP was robustly increased when cells were cultured in low glucose medium, while FGF1-GFP secretion was minimally affected (**Fig. 1k**). No secretion of GFP alone was detected in either high glucose or low glucose condition. Our results indicate that ASRA secretion is mediated in part by secretory autophagy and is stimulated by energy deficiency.

### aP2-*Asra* transgenic mice are hyperphagic and severely obese

To investigate the potential *in vivo* function of ASRA in energy balance, we overexpressed *Asra* in adipose tissue driven by the aP2 promoter. Prior to weaning (∼4 weeks), there was no body weight difference between the aP2-*Asra* transgenic mice and littermate controls (**Extended Data Fig. 2a**). After weaning, male aP2-*Asra* mice displayed a substantial increase in body weight (**Fig. 2a**). At seven weeks, fat mass in aP2-*Asra* mice was doubled and lean mass tended to increase (**Fig. 2b**). At 7 months, male aP2-*Asra* mice became severely obese. They weighed 52.32 ± 0.81 g compared with 36.74 ± 1.0 g for the littermate controls, representing a 42% increase in body weight (**Fig. 2a**), which was accompanied by increased fat mass and liver steatosis (**Extended Data Fig. 2b**). Similarly, female aP2-*Asra* mice had a 40% increase of body weight relative to littermate controls (**Extended Data Fig. 2c and 2d**). Interestingly, both male and female aP2-*Asra* mice had a 10% increase in body length (**Fig. 2c**). Enhanced linear growth has previously been observed in hyperphagic obese mice caused by mutation within genes comprising the LepR signaling pathway and downstream targets, including the LepR Y1138S point mutation (which is incapable of STAT3 signaling)^30^, STAT3 knockout^31^, POMC knockout^32^ and melanocortin-4 receptor (MC4R) knockout^33^, although the exact mechanisms are unclear. The development of obesity in aP2-*Asra* mice was markedly accelerated by a high-fat diet (**Fig. 2d and Extended Data Fig. 2e**). Male aP2-*Asra* mice starting on a high-fat diet at five-week-old for 11 weeks reached a body weight of 55.3 ±1.7 g (**Fig. 2d**), while wild-type mice typically need more than 20 weeks of feeding to reach a body weight of 50 g.

**Figure 2.**
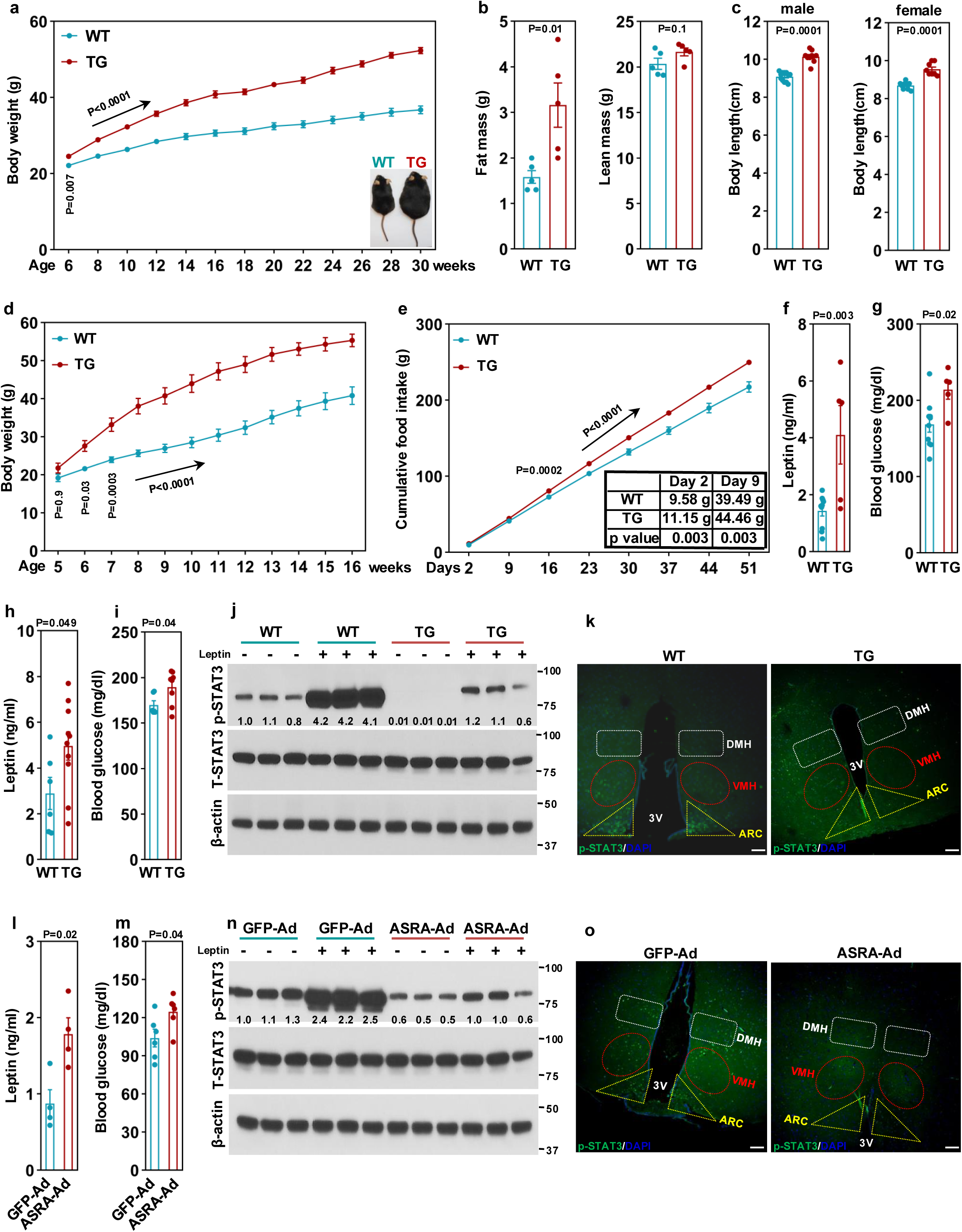
Exogenous expression of ASRA attenuates leptin receptor signaling that leads to hyperphagic obesity. **a,** Body weight of male aP2-*Asra* transgenic mice (TG) (n=12) and littermate controls (n=13) on a chow diet. **b,** Fat mass and lean mass of aP2-*Asra* TG mice (n=5) and littermate controls (n=5) at seven-week-old. **c,** Body length of male (n=9-12) and female (n=8-9) aP2-*Asra* TG mice and littermate controls. **d,** Body weight of male aP2-*Asra* TG mice (n=5) and littermate controls (n=7) on a high fat diet. **e,** Cumulative food intake of aP2-*Asra* TG mice (n=10) and littermate controls (n=11). **f, g,** Circulating leptin (**f**) and glucose (**g**) levels in aP2-*Asra* TG mice (n=5) and littermate controls (n=10) at nine-weeks-old. **h, i,** Circulating leptin (**h**) and glucose (**i**) levels in pair-fed aP2-*Asra* TG mice (n=8) and ad libitum-fed littermate controls (n=6). **j,** Phosphorylation of STAT3 at Tyr705 in hypothalamus of aP2-*Asra* TG mice and littermate controls that were pre-fasted for 3 hr. **k,** Phospho-STAT3 immunostaining in the ARC, VMH and DMH of hypothalamus in aP2-*Asra* TG mice and littermate controls that were fasted for 5 hr followed by leptin injection. Bar=200 μm. **l, m,** Circulating leptin (**l**) (n=4) and glucose levels (**m**) (n=6) of mice injected with either GFP or ASRA adenoviruses. **n,** Phosphorylation of STAT3 in adenovirus-infected mice that were fasted for 3 hr. **o,** Phospho-STAT3 immunostaining in the ARC, VMH and DMH of hypothalamus in adenovirus-infected mice that were fasted for 5 hr followed by leptin injection. Bar=200 μm.

Prior to their development of significant obesity, we examined oxygen consumption, activity, and expression levels of genes for adipose thermogenesis and mitochondrial metabolism in regular diet-fed aP2-*Asra* mice and found no difference compared with littermate controls (**Extended Data Fig. 2f**-2h). Next, mice were individually caged at 5 weeks and their food intake was measured starting at 6 weeks of age for 51 days. We found that the aP2-*Asra* mice were hyperphagic, consuming, on average, 0.70 ± 0.06 g more food per day than littermate controls (**Fig. 2e**). Importantly, hyperphagia was evident at the onset of body weight divergence, as shown by the cumulative food intake at day 2 and day 9 (**Fig. 2e, inset**), suggesting that hyperphagia is the cause, rather than the consequence, of the obesity phenotype. To confirm this, we performed pair-feeding experiments in which the aP2-*Asra* mice were offered an averaged amount of food consumed by the control group on the previous day. Pair-fed aP2-*Asra* mice had similar body weights as control mice fed ad libitum (**Extended Data Fig. 2i**), demonstrating that hyperphagia indeed underlies the obesity phenotype.

### Exogenous expression of ASRA attenuates LepR signaling and causes early onset of leptin resistance and elevated blood glucose

As might be expected, old aP2-*Asra* mice displayed hyperleptinemia, hyperglycemia, and hyperinsulinemia (**Extended Data Fig. 2j**-2l). Interestingly, in young aP2-*Asra* mice, levels of circulating leptin and glucose were significantly elevated as well (**Fig. 2f and 2g**), reflecting leptin resistance. The early onset of leptin resistance promoted us to examine whether this also occurred in pair-fed aP2-*Asra* mice. Indeed, these mice had increased circulating leptin and glucose levels (**Fig. 2h and 2i**), dissociating leptin resistance and elevated glucose from hyperphagia and body weight gain. We next intraperitoneally injected vehicle or leptin into aP2-*Asra* mice and littermate controls and examined STAT3 phosphorylation at Tyr705 in the hypothalamus. Basal STAT3 phosphorylation as well as leptin-induced STAT3 activation was much lower in aP2-*Asra* mice (**Fig. 2j**). Fluorescent immunostaining showed that both the number and intensity of phospho-STAT3 positive neurons in ARC and the ventromedial nucleus (VMH) of the hypothalamus were significantly decreased (**Fig. 2k and Extended Data Fig. 2m and 2n**). These results together suggest that impaired LepR signaling is a primary and early event in aP2-*Asra* mice.

To examine the acute effect of exogenous ASRA expression on LepR signaling, we tail-vein injected wild-type mice with adenoviruses expressing *Asra*, which resulted in production of ASRA protein in the liver and its secretion into circulation (**Extended Data Fig. 2o**). We found that hepatic expression of ASRA similarly increased circulating leptin and glucose levels (**Fig. 2l and 2m**). Furthermore, STAT3 phosphorylation in the hypothalamus (**Fig. 2n**), especially in the ARC and VMH regions (**Fig. 2o and Extended Data Fig. 2p and 2q**), was reduced by hepatic expression of ASRA. These results suggest rapid and direct effects of ASRA on LepR signaling and glucose homeostasis. In aggregate, exogenous expression of ASRA in peripheral tissues can both chronically and acutely attenuate hypothalamic LepR signaling, elevate leptin levels and cause leptin resistance, which, in the aP2-*Asra* mice, are largely responsible for hyperphagia, obesity, and hyperglycemia.

### Adipose-specific and liver-specific ASRA knockout mice have lower food intake and are resistant to high-fat diet-induced obesity (DIO)

We generated conditional *Asra* mice and crossed them with Adiponectin-Cre mice to produce adipose-specific *Asra* knockout (ADKO) mice (**Extended Data Fig. 3a and 3b**). We measured food intake in large cohorts. While food intake of *Asra* ADKO mice in any given day often tended to be lower without statical significance, the cumulative food intake was significantly less compared with that of littermate controls (**Fig. 3a**). The lower food intake did not lead to a lower body weight (**Fig. 3b**), indicating that a compensatory response might have occurred to prevent an unsustainable energy deficit; however, expression of genes for adipose thermogenesis and mitochondrial oxidative metabolism remained unchanged (**Extended Data Fig. 3c and 3d**). We then fed male ADKO mice a high-fat diet. The ADKO mice had a more evident and persistent lower food intake (**Fig. 3c**), and as a result, these mice gained significantly less body weight and had decreased adiposity and liver weight, compared with littermate controls (**Fig. 3d and 3e, and Extended Data Fig. 3e**). Similar results were obtained in female ADKO mice on a high-fat diet (**Extended Data Fig. 3f-3h**).

**Figure 3.**
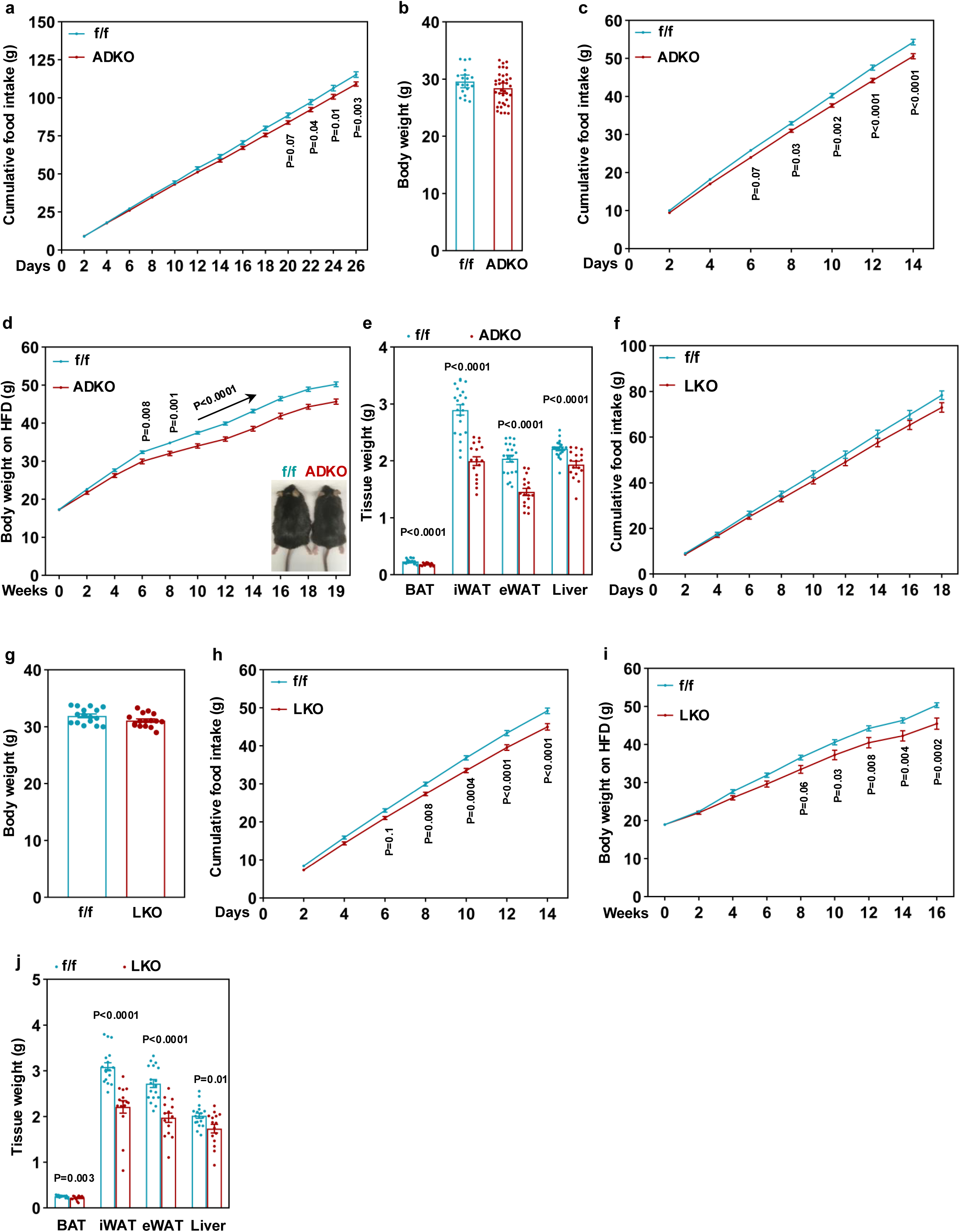
Adipose-specific and liver-specific *Asra* knockout mice have lower food intake and are resistant to DIO. **a,** Cumulative food intake of male *Asra* ADKO mice (n=37) and littermate controls (n=19) on a regular diet. **b,** Body weights of mice in (**a**) on a regular diet. n=19-37/group. **c**, Cumulative food intake of mice in (**a**) on a high fat diet. n=19-37/group. **d,** Body weights of a second cohort of male *Asra* ADKO mice (n=17) and littermate controls (n=20) on a high fat diet. **e,** Tissue weights of mice in (**d**). n=17-20/group. **f,** Cumulative food intake of *Asra* LKO mice (n=16) and littermate controls (n=16) on a regular diet. **g,** Body weights of mice in (**f**) on a regular diet. n=16/group. **h,** Cumulative food intake of mice in (**f**) on a high fat diet. **i,** Body weights of a second cohort of male *Asra* LKO mice (n=15) and littermate controls (n=18) on a high fat diet. **j,** Tissue weights of mice in (**i**). n=15-18/group.

Hepatic *Asra* mRNA level is about 20% of that of WAT. Nonetheless, given the large liver mass, induction of hepatic *Asra* expression by fasting, and the robust capacity of the liver to distribute secreted factors to other tissues^34^, the physiological role of liver-secreted ASRA could be substantial. We obtained *Asra* liver-specific knockout (LKO) mice (**Extended Data Fig. 3i**) by crossing *Asra* conditional mice with Albumin-Cre mice. On normal chow diet, there were no significant differences in food intake and body weight between LKO mice and control mice, although LKO mice had a lower trend (**Fig. 3f and 3g**). On a high-fat diet, similar to the ADKO mice, the LKO mice displayed attenuated food intake, lower body weight gain, and less fat mass and liver mass (**Fig. 3h-3j and Extended Data Fig. 3j**). Our data collectively suggest that endogenous ASRA acts as an orexigenic signal to stimulate food intake and is necessary for normal appetite, and its deficiency in either adipose or liver consequently causes resistance to DIO. Of note, both ADKO and LKO mice maintained normal linear growth (**Extended Data Fig. 3k and 3l**).

### Peripheral ASRA-deficiency sensitizes leptin action and improves glucose homeostasis

To understand the primary mechanism underlying the physiological roles of ASRA, we analyzed leptin signaling in normal chow diet-fed *Asra* ADKO and LKO mice. These mice had a similar body weight as littermate controls; therefore, potential body weight and fat mass differences as confounding factors were eliminated. We found that circulating leptin levels were markedly lower in the KO mice (**Fig. 4a and 4b**), especially in the ADKO mice, which had a 9-fold reduction, indicating increased leptin sensitivity. Thus, in the face of decreased ASRA levels, the *Asra* KO mice accordingly lowered their leptin levels to avoid severe hypophagia and hence maintain metabolic health, underscoring a tight coordination between leptin and ASRA. Interestingly, this adjustment in leptin level was in part mediated at the transcriptional level (**Extended Data Fig. 4a and 4b**), which was also observed in aP2-*Asra* mice (**Extended Data Fig. 4c**). Lower circulating leptin levels coincided with improved glucose homeostasis (**Fig. 4c and 4d**). In further support of the leptin hypersensitivity in *Asra* KO mice, western blot analysis showed that both basal and leptin-stimulated STAT3 phosphorylation in the hypothalamus was higher (**Fig. 4e and Extended Data Fig. 4f**), and fluorescent immunostaining revealed increased leptin-stimulated p-STAT3 signal in the ARC and VMH regions (**Fig. 4f and Extended Data Fig. 4d, 4e, 4g and 4h**). To examine the effect of the heightened leptin sensitivity on appetite, we intraperitoneally injected a low dose of leptin (0.5 mg/kg body weight) every 12 hr and monitored food intake. While, as previously reported^35^, this low-dose leptin injection had little effect on food intake in control mice, it reduced food intake by 20% and 14% in ADKO and LKO mice, respectively (**Fig. 4g and 4h**). Together, our data demonstrate that peripheral ASRA-deficiency potentiates both endogenous and exogenous leptin action, revealing suppression of LepR signaling as a physiological mechanism of ASRA action. SOCS3 and PTP1b are important cell-autonomous negative regulators of LepR-STAT3 signaling axis. It is noteworthy that the phenotypes of *Asra* KO mice largely resemble those of rodents with central deficiency of SOCS3 ^36, 37^ and PTP1b^35, 38^, that is, increased leptin sensitivity, improved glucose homeostasis, reduced food intake and resistance to DIO.

**Figure 4.**
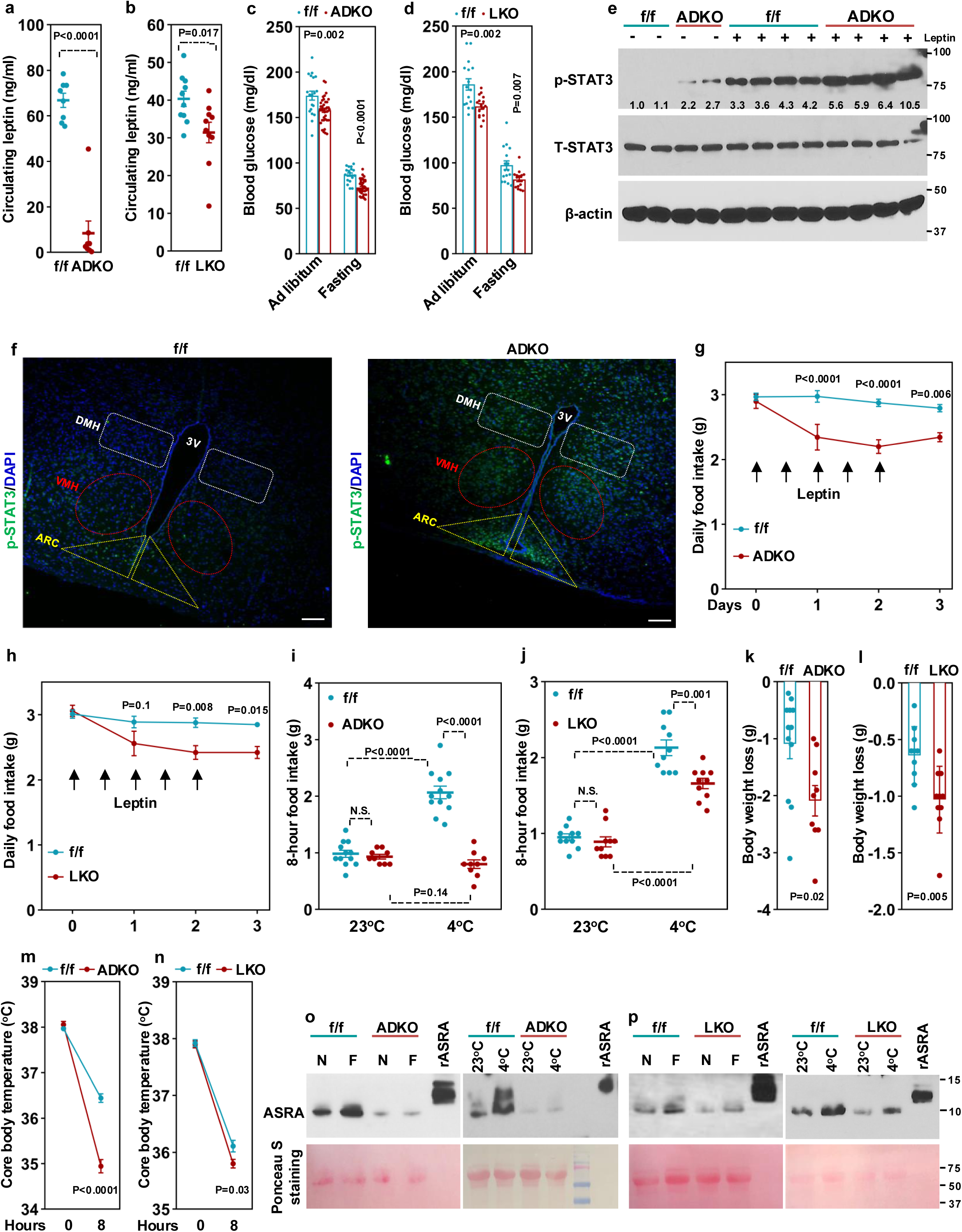
Peripheral ASRA-deficiency sensitizes leptin action and suppresses acute cold-evoked feeding. **a,** Circulating leptin level in five-month-old *Asra* ADKO (n=8) and littermate controls (n=8). **b,** Circulating leptin level in five-month-old *Asra* LKO (n=10) and littermate controls (n=10). **c,** Blood glucose of *Asra* ADKO (n=37) and littermate controls (n=19) fed ad libitum or after 12-hr fasting. **d**, Blood glucose of *Asra* LKO (n=16) and littermate controls (n=16) fed ad libitum or after 12-hr fasting. **e**, Phosphorylation of STAT3 in hypothalamus of mice that were pre-fasted for 3 hr. **f,** Phospho-STAT3 immunostaining in the ARC, VMH and DMH of hypothalamus in ADKO mice and littermate controls that were fasted for 5 hr followed by leptin injection. Bar=200 μm. **g, h,** Daily food intake in male *Asra* ADKO (n=9) and littermate controls (n=12) (**g**) and male LKO (n=10) and littermate controls (n=10) (**h**) that were treated with leptin (0.5 μg/g body weight) twice a day. **i, j,** Food intake in male *Asra* ADKO (n=9) and littermate controls (n=12) (**i**) and male LKO (n=10) and littermate controls (n=10) (**j**) during an 8-hr cold challenge. **k, l,** Body weight loss after cold challenge. **m, n,** Body temperature at 0-hr and 8-hr cold challenge. **o, p,** Three-month-old male *Asra* ADKO mice (**o**) or four-month-old male *Asra* LKO mice (**p**) and littermate controls were either non-fasted (N) or fasted (F) for 12 hr, or were cold challenged for 8 hr. ASRA levels in CSF pooled from 8 mice per group were determined by Western blotting. 0.05 ng of rASRA protein (12 kD) was used as a standard.

### Peripheral ASRA is required for cold-evoked feeding adaptation

Cold stimulates food intake to meet the high energy demand for heat production^14–17^. As ASRA expression is cold-induced in adipose and liver, we investigated whether ASRA is important for cold-stimulated food intake. In an 8 hr cold exposure, control mice consumed 2.07 ± 0.11 g food, more than doubled their food intake at room temperature (**Fig. 4i**). Surprisingly, the ADKO mice consumed 0.8 ± 0.08 g food, similar to their food intake at room temperature (**Fig. 4i**). Consistent with liver being increasingly appreciated as a thermogenesis-responsive organ^39^, the LKO mice also had lower food intake than control mice at cold (**Fig. 4j**), which, together with the complete loss of cold-induced response in the ADKO mice, indicates that there might be a threshold of ASRA level required to enable stimulation of food intake at cold. The KO mice lost more body weight during cold exposure (**Fig. 4k and 4l**), as these mice had to metabolize stored energy to complement inadequate food intake, which was still not sufficient to maintain their core body temperature at the same level as control mice (**Fig. 4m and 4n**). Thus, peripheral ASRA has an essential role in acute cold-evoked feeding, serving as a critical signal that connects thermogenesis to compensatory energy intake.

### ASRA levels in cerebrospinal fluid (CSF) are regulated by fasting and cold

Our results presented so far strongly suggest that peripherally secreted ASRA centrally attenuates LepR signaling. To substantiate our findings, we used western blot analysis along with purified ASRA protein as a standard to estimate endogenous ASRA levels in CSF. We purified His-tagged recombinant ASRA (rASRA) protein from conditioned medium of Expi293F cells transfected with *Asra* expression plasmids. To increase the yield of purified rASRA, the N-terminus of ASRA was fused to a strong classical signal peptide. Of note, the purified rASRA was approximately 12 kD (**Extended Data Fig. 4i**), larger than the expected 8 kD size, likely due to post-translational modification in the classical secretory pathway. ASRA in CSF was readily detected in the wild-type mice, and its level was increased during acute fasting or cold challenge (**Fig. 4o and Fig. 4p**). Importantly, ASRA was markedly decreased in *Asra* ADKO and LKO mice, providing unequivocal evidence that peripherally produced ASRA is a secreted protein and is able to cross the blood-brain barrier. We estimated that ASRA concentration in CSF was in the range of 200-400 ng/L (0.02-0.05 nM) at normal conditions in 3- to 4-month-old wild-type mice and there was a 2-fold induction from the baseline by 12-hr fasting or 8-hr cold challenging (**Extended Data Fig. 4j**). Of note, leptin levels in human CSF are in the range of 0.01-0.02 nM^40^.

### ASRA is a high affinity, orthosteric antagonist of LepR

To further understand how ASRA attenuates LepR signaling, we first examined whether the effect of ASRA can be recapitulated in cell culture. In COS7 cells transfected with the long form of LepR, addition of leptin induced the phosphorylation of STAT3 at Tyr705 and its localization to the nucleus; this activation of STAT3 was abolished by purified rASRA protein in a dose-dependent manner (**Extended Data Fig. 5a and 5b**). Similarly, rASRA dose-dependently suppressed leptin-stimulated activity of a luciferase reporter under control of a STAT3-responsive element (**Fig. 5a**). The half-maximal inhibitory concentration (IC50) of rASRA was 30.00 ± 0.64 nM when 10 nM leptin was used. The effect of rASRA protein was not due to any post-translational modification, as rASRA protein purified from bacteria had a similar effect on LepR signaling (**Extended Data Fig. 5c and 5d**). These results imply that the molecular target of ASRA is within the LepR-STAT3 axis, with LepR a likely target. Importantly, rASRA also decreased the basal activity of LepR (in the absence of leptin), while it had no effect in cells transfected with empty vector (**Fig. 5a**), indicating that ASRA is likely an antagonist/inverse agonist, rather than a partial agonist.

**Figure 5.**
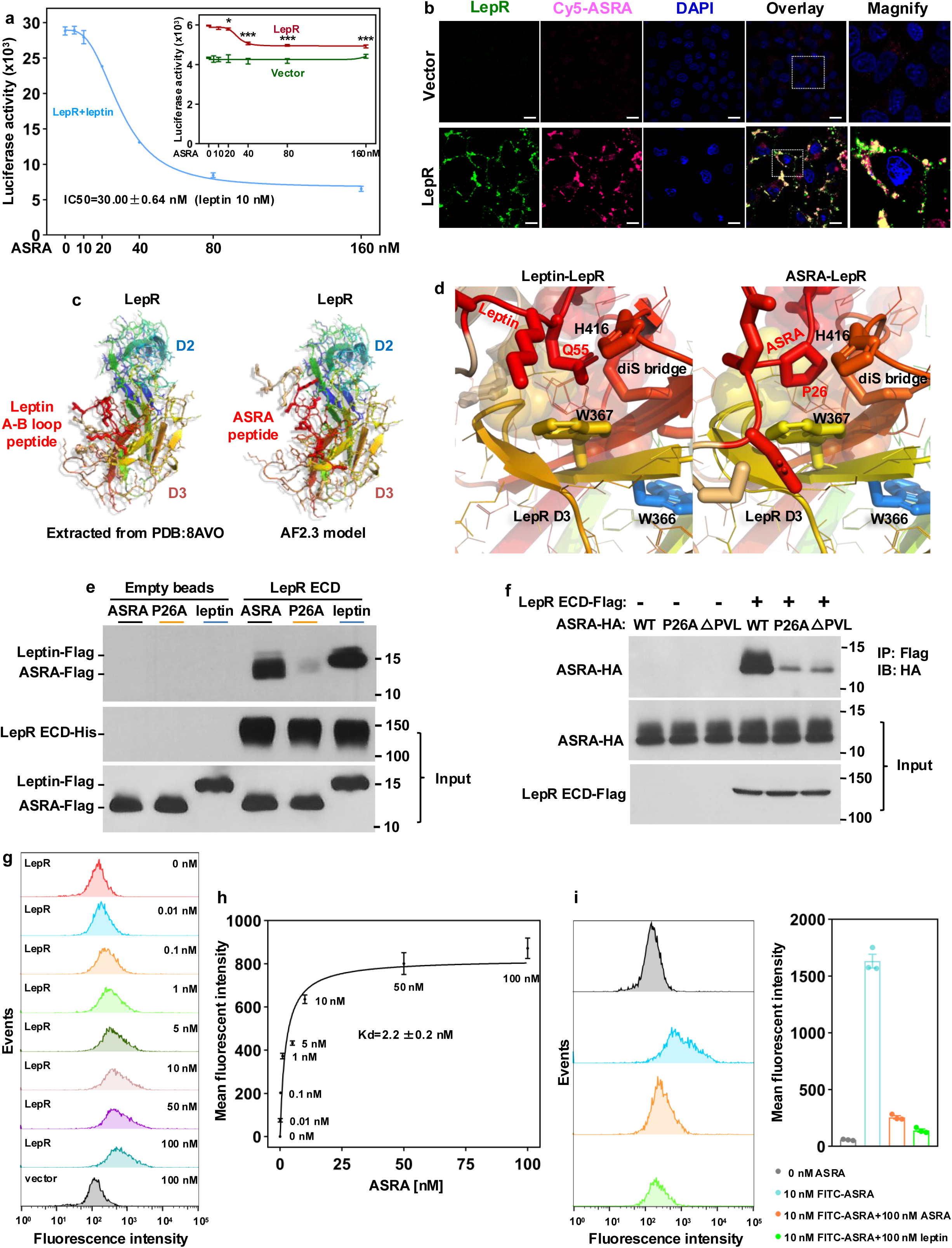
ASRA is a high affinity, orthosteric antagonist of LepR. **a,** HEK293 cells were transfected with indicated plasmids and treated with leptin (10 nM) or vehicle along with rASRA at various concentrations for 24 hours. Luciferase activities were measured (n=3). Note, ASRA also inhibits basal LepR activity, *p<0.05 and ***p<0.001 (versus 0 nM ASRA). **b,** COS7 cells transfected with either vector or LepR plasmids were incubated with Cy5-labeled rASRA protein (300 nM) for 30 minutes. Scale bar = 100 μm. **c,** Comparison of the binding poses of the A-B loop of Leptin docked to LepR D3 domain (left), with the AF2.3-multimer-predicted complex of the ASRA core peptide bound to LepR D3 domain (right). **d,** Key contacts between Leptin’s A-B loop with LepR D3 domain (left), highlighting the sidechain of Gln55 fitting into a pocket defined by Trp367, the Cys413-418 disfulfide bridge, and His416 of LepR. ASRA Pro26 docked to the identical pocket of LepR’s D3 (right). **e,** Equivalent molar concentration of purified ASRA-Flag, ASRA-P26A-Flag, or Leptin-Flag was co-incubated with purified LepR ECD-His or vehicle for 1 hour with a small portion taken as input, followed by precipitation with Ni-NTA beads. After washing, levels of ASRA, leptin and LepR ECD were analyzed by Western blotting. **f,** HEK293 cells were co-transfected with either vector or LepR ECD-Flag plasmids along with HA-tagged wild-type ASRA, ASRA-P26A, or ^26^PVL^28^ deleted ASRA plasmids. Co-immunoprecipitation was performed in conditioned medium. **g,** HEK293 cells were transfected with either vector or LepR plasmids. Cells in suspension were then incubated with FITC-labeled rASRA in PBS and flow cytometry was performed. **h,** Data of average fluorescence intensity per cell were obtained after deduction of background signals and were used to calculate the dissociation constant (Kd) of rASRA binding to LepR (n=3). **i,** HEK293 cells transfected with LepR plasmids were co-incubated with 10 nM FITC-labeled ASRA and 100 nM unlabeled ASRA or leptin. Flow cytometry was performed and average fluorescence intensity per cell was quantified (n=3).

To examine whether ASRA interacts with LepR, HEK293 cells were co-transfected with plasmids expressing the extracellular domain (ECD) of LepR and plasmids expressing either ASRA or leptin. ASRA was co-immunoprecipitated from the conditioned medium with LepR ECD at a similar efficiency as leptin (**Extended Data Fig. 5e**). Moreover, ASRA did not associate with leptin (**Extended Data Fig. 5f**), excluding the possibility that ASRA suppresses LepR signaling by sequestering leptin. Next, cells transfected with full-length LepR were incubated with Cy5-labeled rASRA protein. We found that rASRA not only bound to the cell surface in a LepR-dependent manner but also completely colocalized with LepR (**Fig. 5b**).

The above data suggest that ASRA binds to LepR ECD. How might this occur at a structural level? Leptin binds to LepR at two distinct sites that are both essential for agonistic signaling^41–43^: the high affinity Site 2 within the second cytokine receptor homology region (CRH2) (D4 and D5 domains), and the low affinity Site 3 within the immunoglobulin (Ig)-like domain (D3 domain). While ASRA is largely unstructured, AlphaFold2-multimer (implemented on ColabFold)^44–46^ detects a strongly focused coevolutionary signal between ASRA and LepR chains, that drives the template-free modeling of a nine-residue ASRA peptide ^23^DEAPVLNDK^31^, which is highly conserved among different species (**Extended Data Fig. 1b**), docked to the Ig D3 domain of LepR, recapitulating the binding of the Site 3 A-B loop peptide of leptin^42, 43^ (**Fig. 5c**). The ASRA peptide-LepR D3 complex model, which is reaffirmed by the more recent AlphaFold3 algorithm^47^, reveals several key Site 3-like contacts (**Fig. 5d**); notably, the ring of ASRA Pro26 packs into a pocket against the conserved LepR Trp367 aromatic ring and the adjacent Cys413-Cys418 disulfide bridge--overlapping the Site 3 interaction observed for leptin’s core Gln55 with LepR^42, 43^.

To test this structure model, we purified His-tagged LepR ECD and Flag-tagged ASRA and leptin and performed in vitro binding assays. Similar to the results observed in **Extended Data Fig. 5e**, wild-type rASRA protein was associated with LepR ECD at a similar level to leptin, indicating a similar affinity. Importantly, the rASRA P26A mutant protein largely lost its binding to LepR ECD (**Fig. 5e**), which was also observed in co-immunoprecipitation experiments using conditioned medium (**Fig. 5f**). These results demonstrate a direct interaction between ASRA and LepR ECD and support the predicted structural model. While our data suggest a strong binding of ASRA to Site 3, we cannot rule out the possibility that other regions of the LepR ECD, e.g., Site 2, might be involved as well, as the ASRA P26A mutant retained residual binding activity (**Fig. 5e and 5f**) and ASRA binding was weakened by mutations in the CRH2 domain of the LepR **(Extended Data Fig. 5g**).

Finally, we used flow cytometry to quantitatively analyze the binding of rASRA to full-length LepR on cell surface. FITC-labeled ASRA bound to LepR in a concentration-dependent manner and no binding was detected when LepR was absent. Evident binding was observed even at 0.01 nM ASRA and was largely saturated at 10 nM ASRA (**Fig. 5g and 5h**), which was competed away by 10-fold excess unlabeled ASRA or leptin (**Fig. 5i**). Consistent with their strong interaction observed in **Fig. 5e and Extended Data Fig. 5e**, rASRA bound to LepR with a dissociation constant (Kd) of 2.2 ± 0.2 nM (**Fig. 5h**), while the Kd for leptin is in the range of 0.3-1.2 nM^48–50^. These data together strongly suggest that ASRA is a high affinity, orthosteric antagonist of LepR.

### rASRA protein stimulates food intake and elevates blood glucose in a LepR signaling-dependent manner

rASRA injected intraperitoneally was detected in CSF (**Extended Data Fig. 6a**). Moreover, injected Cy5-labeled rASRA bound to the ARC of the hypothalamus in a pattern that was highly overlapped with LepR (**Extended Data Fig. 6b**). To determine the *in vivo* effects of rASRA, we intraperitoneally injected wild-type rASRA or vehicle into individually caged wild-type lean mice and food intake was measured every 24 hr. A single dose of rASRA significantly stimulated food intake in wild-type mice (**Fig. 6a**, Day 1, p=0.0003). rASRA purified from bacteria had a similar orexigenic effect, which can be observed at the very early phase of rASRA injection (**Fig. 6b**), resembling the acute effect of Ghrelin^51^ but distinct from the delayed effect of Asprosin^7^. We then injected rASRA once a day for 9 days. rASRA-treated wild-type mice continued to have a higher food consumption (**Fig. 6a**). No body weight gain was observed (**Fig. 6c**), likely due to the short-term treatment and individually-caging environment, which augments energy expenditure. Remarkably, this treatment elicited a more than 10-fold increase in leptin level (**Fig. 6d**) concomitant with an elevated glucose level (**Fig. 6e**). Thus, rASRA-treated mice mounted a compensatory response attempting to avoid overfeeding, which demonstrates both antagonism and coordination between leptin and ASRA. Despite hyperleptinemia, the rASRA-treated mice were highly leptin-resistant with a lower basal STAT3 activity in the hypothalamus (**Fig. 6f**), consistent with increased food intake. In contrast, treatment of wild-type mice with rASRA P26A mutant protein had no effects on food intake, circulating leptin and glucose levels, or hypothalamic STAT3 phosphorylation (**Fig. 6g-6j**). We also treated ob/ob mice with wild-type rASRA and found that neither food intake nor blood glucose was affected (**Fig. 6k and Fig. 6l**). Thus, ASRA-stimulated food intake requires its interaction with LepR and a functional leptin signaling pathway. Next, we rescued leptin signaling in ob/ob mice by injection of leptin. While exogenous leptin stimulated STAT3 phosphorylation in hypothalamus and effectively suppressed food intake in ob/ob mice as expected, these leptin-induced effects were markedly attenuated by co-injection of rASRA (**Fig. 6m and 6n**). These data together provide strong *in vivo* evidence that ASRA directly antagonizes LepR signaling to stimulate appetite.

**Figure 6.**
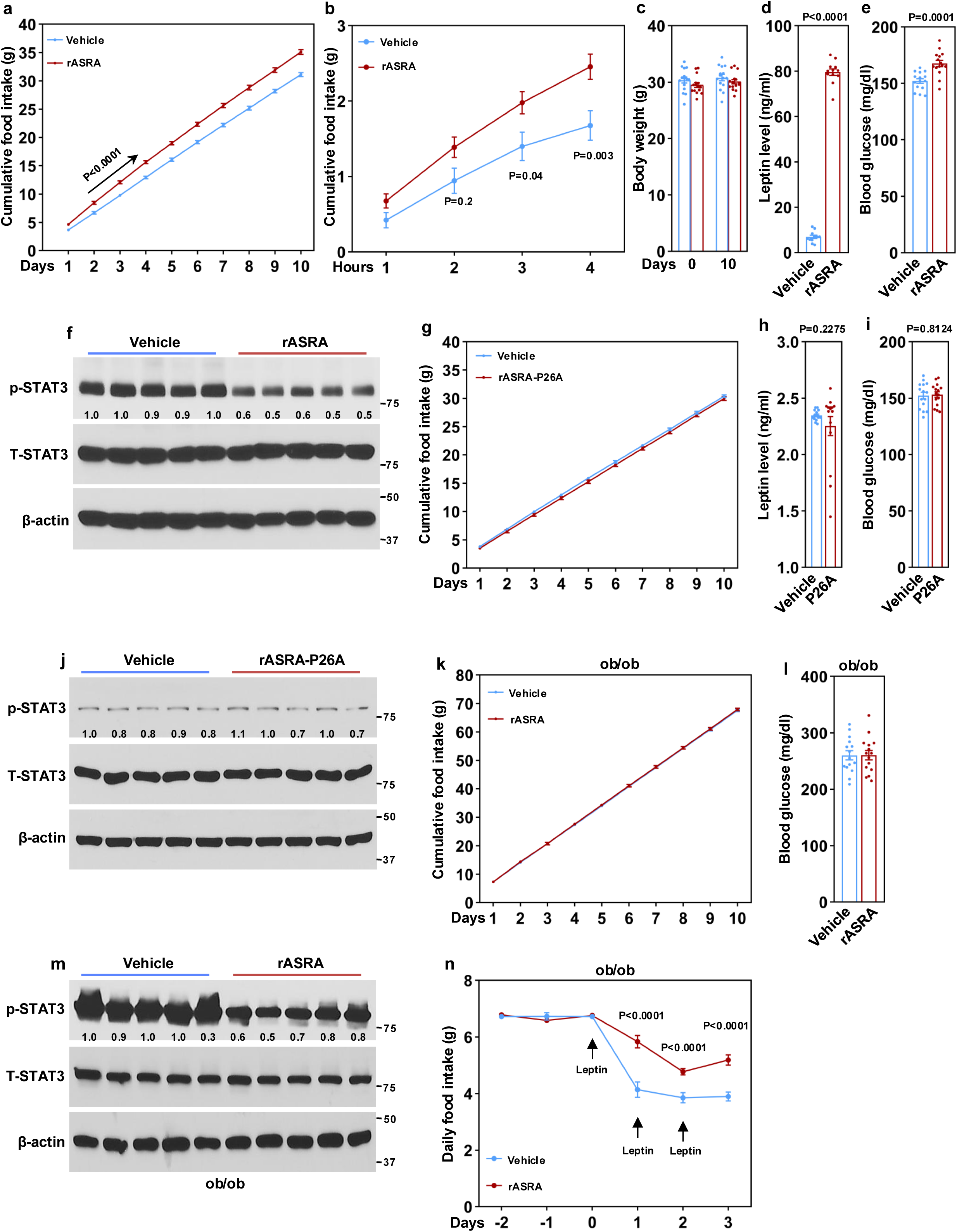
rASRA protein induces hyperleptinemia, and stimulates food intake in a leptin receptor signaling-dependent manner. **a,** Three-month-old male mice were daily injected with rASRA protein (65 μg per mouse per day), and cumulative food intake was measured. n=15 per group. **b,** A single injection (65 μg) of rASRA purified from bacteria was ip injected, and cumulative food intake was measured. n=9 per group. **c,** Body weight of mice in (**a**). n=15 per group. **d,** Leptin levels of mice in (**a**) at day 10. Twelve serum samples per group were randomly picked. **e,** Blood glucose of mice in (**a**) at day 10. n=15 per group. **f,** Basal phospho-STAT3 in the hypothalamus of mice in (**a**) without fasting. **g,** Three-month-old male mice were daily injected with rASRA-P26A protein (65 μg per mouse per day), and cumulative food intake was measured. n=15 per group. **h,** Leptin levels were measured at day 10 from mice in (**g**). n=15 per group. **i,** Blood glucose of mice in (**g**) was measured at day 10. n=15 per group. **j,** Basal phospho-STAT3 in the hypothalamus of mice in (**g**) without fasting. **k,** Three-month-old male ob/ob mice were daily injected with rASRA protein (100 μg per mouse per day), and cumulative food intake was measured. n=15 per group. **l,** Blood glucose of mice in (**k**) was measured at day 10. n=15 per group. **m, n,** Fourteen-week-old male ob/ob mice were treated with rASRA (100 μg per mouse per day) or vehicle along with leptin (2 μg/g body weight per day) for 3 days. Phosphorylation of STAT3 in hypothalamus (**m**) and food intake (**n**) were measured. n=15 per group.

## Discussion

In this work, we identified ASRA, produced by adipose and liver, as a potent orexigenic peptide that centrally suppresses LepR signaling and regulates appetite and glucose metabolism. Intraperitoneal administration of wild-type rASRA protein, but not that of rASRA P26A mutant protein defective in LepR-binding, causes hyperleptinemia, suppresses LepR signaling, and stimulates food intake. Mice with chronic overexpression of ASRA in adipose tissue are hyperphagic, severely obese, hyperleptinemic, and hyperglycemic. On the other hand, ablation of ASRA in either fat or liver sensitizes leptin action, improves glucose homeostasis, reduces food intake, and leads to resistance to DIO. Furthermore, we found that peripheral ASRA is indispensable for acute cold-stimulated food intake, a process that has been poorly understood. Thus, ASRA serves as a built-in, endocrine rheostat robustly counteracting LepR signaling with both physiological and pathophysiological significance. Interestingly, this ASRA antagonism is intrinsically coordinated with leptin, as in response to ASRA level, a feedback loop regulating leptin level, independent of body weight and adiposity, is rapidly activated to avoid excessive overfeeding or severe hypophagia, underscoring that ASRA and leptin function as an integrated system.

There are precedents that orexigenic and anorexigenic peptides with distinct amino acid sequences can impinge upon the same receptor^5, 52^. Our data show that ASRA stimulates appetite by acting as a high affinity, orthosteric antagonist of LepR, despite bearing no sequence similarity with leptin. While recognition of the leptin A-B loop by the Ig D3 domain of LepR is considered a low affinity interaction, ASRA, through a conserved core peptide, appears to competitively bind to this same site, but with a much higher affinity; further structural studies are needed to investigate how this high-affinity binding is achieved. The mechanistic action of ASRA distinguishes it from the other two known peripherally produced orexigenic peptides, Ghrelin and Asprosin^6, 7^, which function independently of LepR. In addition to the endocrine antagonism we identified here, the LepR-STAT3 signaling axis is also negatively feedback-modulated by cell-autonomous regulators, emphasizing the need for tight control of this crucial signaling pathway by both peripheral and central regulators at different molecular levels. These negative regulatory mechanisms may have reinforcing or complementary roles in modulating LepR signaling under different physiological and pathophysiological circumstances. Adding another layer of regulation, a recent study shows that an HDAC6-regulated, adipose-secreted protein, though it remains to be identified, potentiates leptin action^53^.

Given the small mass of BAT, despite its enrichment of ASRA expression, the endogenous endocrine action of ASRA is conceivably attributed primarily to WAT- and liver-derived ASRA. ASRA mRNA expression in both adipose and liver and ASRA protein level in CSF are increased by fasting. In contrast, leptin mRNA expression in adipose^54, 55^ and serum leptin level^56–58^ fall during fasting. This opposing regulation in response to fasting aligns well with the action of ASRA, especially given observations that a substantial level (e.g., 20-30%) of circulating leptin is still present even after a long fasting period in rodents^56–58^. Thus, the ASRA antagonism appears to constitute a timely signaling modality to ensure sufficient feeding response, allowing a quick restoration of energy balance. In this regard, it is interesting to note that ASRA associates with autophagy vesicles, and its secretion is enhanced in response to energy deficiency, which may represent an intriguing, previously unappreciated link between peripheral cellular autophagy and hypothalamic regulation of appetite.

Increased energy expenditure in the cold elicits a compensatory hyperphagic response to maintain adiposity and body weight, and a peripheral feedback signal to the hypothalamus has been postulated^17^. Similar to fasting, ASRA mRNA expression in adipose and liver and ASRA protein level in CSF are increased by cold, whereas leptin expression in adipose^59^ and serum leptin level^58^ fall. Remarkably, peripheral ASRA deficiency in either adipose or liver profoundly impairs cold-stimulated food intake, with the ADKO mice being completely unresponsive. It is also interesting to note that aP2-Hlx transgenic mice display a remarkably heightened adipose thermogenesis, and as a result, are severely hyperphagic associated with an increase of adipose ASRA expression^18^. Our results collectively suggest that peripheral ASRA transmits a signal of energy deficiency to the hypothalamus to promote food intake. From an evolutionary perspective, this ASRA antagonism may allow a survival advantage by stimulating food-seeking to defend hunger and cold.

At the other end of the spectrum, however, ASRA antagonism may be one of the pathogenic mechanisms underlying obesity-driven leptin resistance. Studies have suggested that leptin resistance involves an overactivation of cell-autonomous negative regulators. It would be intuitive that a humoral factor, perhaps correlated with obesity or adiposity, also plays a part. Given the expansion of adipose tissue and elevated ASRA expression in liver in DIO mice, total ASRA level is expected to markedly increase in circulation and CSF. According to our data and as would be expected for a LepR antagonist, such an increase of circulating ASRA would not only exacerbate leptin resistance but also drive elevated leptin levels, which may lead to a vicious cycle in the state of obesity. We thus propose that ASRA is an important contributor to the development of leptin resistance and progression toward severe obesity. While our current study has provided important evidence suggesting this is plausible, future investigation using ASRA neutralizing antibodies or simultaneously deleting ASRA in both adipose and liver in leptin resistance-established DIO mice should further allow us to rigorously address its role.

In summary, our studies reveal a previously unknown mechanism of hormonal control of appetite in which a peripherally secreted protein ASRA serves as a high-affinity antagonist of LepR in the hypothalamus to attenuate leptin signaling, which may have important implications for our understanding of leptin resistance. ASRA has the potential to be a therapeutic target for obesity, diabetes, Prader-Willi syndrome, anorexia, and cancer cachexia.

## Acknowledgments

We thank the Transgenic Animal Modeling Core at University of Massachusetts Chan Medical School for the generation of ASRA transgenic mice, the Morphological Core for help on histology, and the Metabolic Core for metabolic cage studies. We thank Dr. Eric Baehrecke for providing RFP-LC3, mCherry-ER and RFP-Golgi plasmids, and Dr. Jan Tavernier for providing pXP2d2-rPAPI-luciferase plasmid. We thank Drs. Mike Czech and Eric Baehrecke for comments on the manuscript. This work was supported by NIH R01DK116872 and R01DK115918 (to Y.-X.W.). P.P.L. and S.A.W. were supported in part by NIH 1UH3TR002668 and 1R01HL150669.

## Author Contributions

L.H., P.P.L, Y.D., and D.P. designed and performed experiments, and analyzed data. J.F.B. performed structural prediction of ASRA-LepR complex. P.P.L. performed bioinformatics analysis. Y.-X.W. designed experiments and analyzed data. S.A.W. analyzed data. Q.C., A.L. and V.K. contributed technical assistance. L.H. and Y.-X.W. wrote the manuscript with contributions from J.F.B. and Y.D. S.A.W. edited the manuscript.

## Methods

### Identification of ASRA

Genes that are abundantly expressed in BAT (FPKM > 30), and have > 2-fold and > 10-fold expression levels in BAT relative to the levels in epididymal WAT and soleus muscle, respectively, were selected from gene expression dataset GSE56367 ^19^, and were further screened using dataset GSE7623 ^20^ for the genes within this subset that are induced by fasting. Overlapping genes from the above analyses were then predicted for secretion potential using SignalP and SecretomeP (www.cbs.dtu.dk/services)^21, 22^, which resulted in the identification of 1190005I06RIK and its human orthologue C16orf74 (ASRA).

### Mice

C57BL/6J wild-type mice (Stock No. 000664) and B6.Cg-*Lep^ob^*/J mice (Stock No. 000632) were purchased from Jackson Laboratory. ASRA transgenic mice were generated at the core facility of UMASS Chan Medical School. Briefly, the ASRA cDNA was inserted downstream to the 5.4 kb aP2 promoter. The transgenic DNA fragment was purified using gel electrophoresis and then injected into fertilized embryos obtained from C57BL/6J×SJL hybrid mice. Transgenic lines were bred with C57BL/6J mice for a minimum of three generations. *Asra* conditional knockout mice, in which exon 3 and exon 4 were flanked by two loxP sites, were generated by Biocytogen using CRISPR/Cas9 technology. The resulting floxed mice were then crossed with Adiponectin-Cre mice^60^ or Alb-Cre mice^61^ (Stock No. 003574, Jackson lab) to delete the last 67 amino acid residues of ASRA in adipose tissue or liver, respectively. Mice were maintained under a 12 hours light/12 hours dark cycle at 23°C with free access to food and water, unless otherwise indicated. Mice were fed either a regular diet containing 4% (w/w) fat or a high-fat diet (Bioserv, cat#S3282) containing 36% (w/w) fat. Gender-matched littermate controls were used in all experiments, and their ages were indicated accordingly. All animal studies were conducted in accordance with the guidelines approved by the Institutional Animal Care and Use Committee (IACUC) at the University of Massachusetts Medical School.

### Plasmids

Mouse *Asra* (encoding 76 amino acid residues) and *Fgf1* cDNA were generated by PCR. ASRA-GFP and FGF1-GFP were constructed by fusing GFP to the C-terminus of ASRA or FGF1 in a mammalian expression vector. The long form of human *LepR (LepRb)* cDNA was obtained from a commercial source, and point mutations were generated by PCR; all were verified by sequencing. RFP-LC3, mCherry-ER and RFP-Golgi plasmids were kindly provided by Dr. Eric Baehrecke’s laboratory. The STAT3 luciferase reporter plasmid pXP2d2-rPAPI-luciferase^62^ was kindly provided by Dr. Jan Tavernier’s laboratory.

### Hepatic overexpression of ASRA through adenovirus infection

Adenoviral ASRA overexpression plasmid was generated using the AdEasy-1 system^63^. The plasmid was transfected into the AD-293 cells, and the adenoviruses were purified through cesium chloride ultracentrifugation. The viral titers were determined by counting GFP-positive HEK293 cells infected with the viruses. Wild-type mice were tail-vein infused with GFP or ASRA adenoviruses at 1 × 10^10^ moi/mouse. One week after infection, serum and hypothalamus were collected.

### Food intake

The mice were individually housed for at least three days with ad libitum access to either a regular chow or high-fat diet prior to commencing the food intake measurements. Food intake was measured daily or at indicated time intervals. To examine the effect of leptin on food intake, *Asra* knockout mice and littermate controls were intraperitoneally (ip) injected with leptin at 0.5 μg/g body weight twice a day (at 8 am and 8 pm). To measure cold-invoked feeding, *Asra* knockout mice and littermate controls were housed in 4°C cold room for 8 hr from 9 am to 5 pm. To perform pair-feeding experiments, mice were housed individually for a period of 5 days, during which they had unrestricted access to regular chow. Subsequently, the aP2-*Asra* mice were provided with an amount of food equivalent to the average consumption of wild-type mice over the preceding 24 hour period, and body weight was monitored.

### Expression and purification of recombinant proteins

To aid the purification, wild-type *Asra* cDNA, *Asra*-P26A cDNA and *LepR*-ECD cDNA were cloned into pHL-sec vector (Addgene)^64^ to add a signal peptide at N-terminus and a 6xHis tag at C-terminus. The expression plasmids, after removal of endotoxin, were transiently transfected into mammalian Expi293F cells (ThermoFisher, cat#A14527) cultured in suspension. Medium was collected, loaded onto a Ni-NTA Affinity column, and washed with Ni-NTA Binding/Wash Buffer (1mM TCEP, 20 mM TRIS (pH 7.5), 40 mM Imidazole, 1000 mM NaCl). Protein was eluted with Elution buffer (8% Glycerol (% V/V), 1mM TCEP, 20mM TRIS (pH 7.5), 500 mM Imidazole, 500 mM NaCl), and then concentrated and buffer-exchanged into PBS via an Amicon Ultra-15 Centrifugal Filter with a 3 kDa cut-off (Millipore, cat#UFC900324). Purified protein was subjected to sterile filtration utilizing a COSTAR 0.22 μm Spin-X Centrifuge Tube Filter. Concentrations were measured by BCA assay. Protein was divided into smaller portions and stored at −80°C, with a maximum of three freeze-thaw cycles. For *in vivo* use of rASRA, endotoxin level (0.01 EU/μg) was measured with a commercial kit (Thermo Scientific, cat#88282).

To produce ASRA-Flag and Leptin-Flag proteins, wild-type *Asra* cDNA, *Asra*-P26A cDNA, and *LepR*-ECD cDNA were cloned into the pHL-sec vector with a Flag tag at the C-terminus. The plasmids were transiently transfected into Expi293F cells, and the medium was harvested after 72 hours. The medium was then loaded onto an ANTI-FLAG® M2 affinity gel (Sigma, cat#A2220) chromatography column and washed with 1X wash buffer (50 mM Tris-HCl, pH 7.4, 150 mM NaCl). The Flag-tagged proteins were eluted using 3X FLAG^®^ peptide (Sigma, cat#F4799) at a concentration of 200 ng/µL in 1X wash buffer. Finally, the proteins were concentrated and buffer-exchanged into PBS using an Amicon Ultra-15 Centrifugal Filter with a 3 kDa cut-off.

To purify rASRA from bacteria, *Asra* cDNA with a 6xHis tag was cloned into the pET23b vector. The *Asra* bacterial expression plasmid was then transformed into BL21 (DE3) cells. rASRA protein production was induced by addition of 0.5 mM IPTG, followed by overnight incubation at 18°C. Following cell lysis, cleared cell extracts were loaded onto a Ni-NTA Affinity column, extensively washed, and eluted. rASRA protein was then concentrated and buffer-exchanged into 20 mM Tris-HCl and 150 mM NaCl. Any residual endotoxin was removed using the Pierce High Capacity Endotoxin Removal Spin Column (ThermoFisher, cat#88274). The endotoxin level was 0.001 EU/μg.

### *In vivo* administration of rASRA protein

rASRA and rASRA-P26A proteins purified from Expi293F cells were ip injected into wild-type mice or ob/ob mice at 65 μg/mouse or 100 μg/mouse per day, respectively. Leptin at 2 μg/g body weight per day was ip injected into ob/ob mice in co-injection experiments. In separate experiments, a single dose of rASRA protein purified from bacteria was ip injected into wild-type mice at 65 μg/mouse body weight. Cumulative food intake was measured at indicated time points.

### STAT3 activation

To measure STAT3 activation in the hypothalamus, mice were ip injected with leptin at 1 μg/g body weight. After 45 min, mice were euthanized and hypothalamuses were obtained. Activated STAT3 (phospho-Tyr705) (Cell Signaling Technology, cat#9145) and total STAT3 (Cell Signaling Technology, cat#9139) were determined by western blot analysis. For immunofluorescence staining, hypothalamic sections that were fixed in formalin and embedded in paraffin were subjected to deparaffinization and rehydration using graded ethanol solutions. Following pre-incubation with a blocking buffer (PBS containing 5% normal goat serum and 0.3% Triton X-100) at room temperature for 60 minutes, the slides were incubated overnight at 4°C with a 1:100 dilution of phospho-STAT3-Tyr705 antibody (Cell Signaling Technology, cat#9145) in blocking buffer. Subsequently, the slides were washed and incubated with Alexa Fluor 488-conjugated secondary antibody (ThermoFisher, cat#A-11008) and DAPI (Sigma, cat#D9542) at room temperature for 2 hours.

To determine STAT3 activation in cell culture, COS7 cells were transfected with vector or human LepR expression plasmids using jetOPTIMUS DNA transfection reagents (Illkirch, France, cat#101000006), following the manufacturer’s instructions. After a 48-hour incubation period, cells, serum-starved for 6 hours, were pre-incubated with rASRA protein (300 nM) for 30 min and leptin (100 nM) was then added. After 30 min, COS7 cells were fixed with 3.7% formalin at 37°C for 15 minutes and then rinsed twice with PBS. The cells were then permeabilized with ice-cold 100% methanol at −20°C for 10 minutes and washed twice with PBS. Subsequently, the cells were incubated in blocking buffer (PBS containing 5% normal goat serum and 0.3% Triton X-100) at room temperature for 60 minutes. The cells were then incubated overnight at 4°C with a 1:100 dilution of phospho-STAT3-Tyr705 antibody and a 1:200 dilution of β-actin (Santa Cruz Biotechnology, cat#sc-47778) in blocking buffer. The cells were gently washed twice with PBS and incubated with Alexa Fluor 488-conjugated (ThermoFisher, cat#A-11008) and Alexa Fluor 594-conjugated secondary antibody (ThermoFisher, cat#A-11032) for 2 hours at room temperature. Finally, the cells were incubated with DAPI (cat# D9542, Sigma) at room temperature for 30 minutes. Images were captured using an inverted Nikon Eclipse Ti2 confocal microscope (Nikon Instruments/Nikon Corp) and were processed using the same settings.

### Luciferase Reporter Assay

HEK293T cells were cultured in 48-well plates and were transfected with a STAT3 luciferase reporter plasmid (pXP2d2-rPAPI-luciferase) along with human LepR. A vector plasmid was included to maintain the same amount of plasmid transfected in each well. Cells were then pre-treated with indicated concentrations of rASRA for 60 min and 10 nM leptin or vehicle was added and incubated for overnight. Luciferase activity was measured by chemiluminescence with a Synergy H4 Hybrid microplate reader (BioTek, Winooski, VT) using Gen5 software. IC50 was calculated with four-parameter logistic curve fit.

### ASRA and LepR colocalization

rASRA protein purified from Expi293F cells was labeled with Cyanine5 maleimide (Cy5) as per the manufacturer’s instructions (cat#43080, Lumiprobe). COS7 cells were transfected with either vector, human LepR, or LepR mutants. After a 48-hour incubation period, the cells were gently washed with PBS, and Cy5-rASRA (300 nM) was added to the cells in FBS-free medium for 30 minutes at 37°C. The cells were then gently washed twice with PBS and fixed with 3.7% formalin at 37°C for 15 minutes without permeabilization. Next, the cells were incubated in blocking buffer (PBS containing 5% normal goat serum) at room temperature for 60 minutes. The cells were then incubated overnight at 4°C with a 1:100 dilution of LepR antibody (ThermoFisher Scientific, cat#PA1-053) in blocking buffer. After incubation, the cells were gently washed twice with PBS and incubated with Alexa Fluor 488-conjugated goat anti-rabbit IgG (ThermoFisher, cat#A-11008) for 2 hours at room temperature. Finally, the cells were incubated with DAPI (Sigma, cat#D9542) at room temperature for 30 minutes. Images were captured using an inverted Nikon Eclipse Ti2 confocal microscope (Nikon Instruments/Nikon Corp) and were processed using the same settings.

### Structure Modeling

The sequence of human ASRA is disordered in the baseline AlphaFold database (https://alphafold.ebi.ac.uk/entry/Q96GX8). Using the multimer-capable form of AF2.3 with ColabFold 1.5.5 ^44–46^, we narrowed the likely interaction site of ASRA with LepR Ig D3 domain to a core peptide segment of residues 21-32. The cryoEM structure of the heterotrimeric 3:3 Leptin-LepR ectodomain complex (PDB:8AVO)^42^ was superposed with the AF2 prediction and visualized using PyMOL 2.5 (www.pymol.org). The most recent AF3 algorithm^47^, used through its DeepMind portal (https://alphafoldserver.com), corroborated the AF2-multimer results for ASRA binding to LepR.

### Protein binding assays

In co-immunoprecipitation assays, HEK293T cells were co-transfected with LepR-ECD-Flag or vector along with either ASRA-HA, ASRA-P26A-HA, ASRA-△PVL-HA, or leptin-HA, which were all expressed from pHL-sec vector (Addgene)^64^ that contains a signal peptide. After 24 hours, cells were cultured in conditioned medium for 24 hours. The conditioned medium was collected, and immunoprecipitation was performed with anti-Flag M2 affinity gel (Sigma, cat# A2220) for 3 hours at 4°C. The beads were washed four times with buffer [150 mM NaCl, 50 mM Tris (pH 7.5), 0.1% NP-40, 3% glycerol] containing PMSF and protease inhibitor. Immunoprecipitates were analyzed by Western blotting with an anti-HA antibody (Roche, cat#12013819001). To determine whether ASRA associates with leptin, HEK293 cells were co-transfected with leptin-Flag or vector along with ASRA-HA and co-immunoprecipitation was performed in conditioned medium as above.

In binding assays with purified proteins, 0.3 μg purified ASRA-Flag, 0.3 μg ASRA-P26A, or 0.4 μg Leptin-Flag was co-incubated with 5 μg purified LepR ECD-His or vehicle in 1 ml buffer [150 mM NaCl, 50 mM Tris (pH 7.5), 0.1% NP-40, 3% glycerol] for 1 hour at 4°C, followed by addition of 25 µl HisPur™ Ni-NTA beads (Thermo Scientific, cat# 88221) and incubated for 1 hour. The beads were then washed four times and resuspended in 100 µl 1x SDS sample buffer. Levels of ASRA, leptin and LepR ECD were analyzed by Western blotting.

### Flow cytometry

rASRA was conjugated to FITC and excess FITC was removed using the Pierce™ FITC Antibody Labeling Kit (Pierce, cat# 53027) according to the manufacturer’s protocol. We used flow cytometry to measure binding affinity as previously descried^65–67^. HEK293T cells were transfected with either a vector or human LepR. After 48 hours, the cells were dissociated using trypsin/EDTA saline and resuspended in PBS at a density of 300 cells/µl. 100 µl cells were mixed with 100 µl FITC-labeled rASRA in PBS at various final concentrations (0, 0.01, 0.1, 1, 5, 10, 50, and 100 nM) and a final cell density of 1.5×10^5^ cells/ml. Cells were incubated for 2 hours at 4°C in the dark with slow shaking. The 2-hour incubation, which allows to reach a binding equilibrium, was pre-determined. Following incubation, flow cytometry was performed using a Guava easyCyte™ Benchtop Flow Cytometer. The data were analyzed with FlowJo v10.8.1 software. No specific-binding was detected in cells transfected with vector. Data of average fluorescent intensity per cell were obtained after deduction of background signals and were used to fit the built-in one-site specific binding model in Prism 8.1 software to obtain the dissociation constant. For the competition experiments, HEK293T cells transfected with human LepR were co-incubated with 10 nM FITC-labeled ASRA and 100 nM unlabeled ASRA or leptin for 2 hours at 4°C.

### Estimation of ASRA levels in cerebrospinal fluid (CSF)

The *Asra* ADKO, LKO and littermate control mice were fasted overnight or placed in 4°C cold room for 8 h. The collection of mouse CSF was carried out following previously described methods^68^. Briefly, a glass capillary was positioned behind the mouse’s head at a 30-45° angle, ensuring that the sharp end was positioned just beyond the membrane. A volume of 100-200 µL of fluid was drawn using the syringe to establish negative pressure and remove contaminants. The membrane was approached using a micromanipulator, with care taken to avoid puncturing it while sensing resistance. Subsequently, the capillary tube was tapped through the membrane using the micromanipulator, while observing under a microscope as CSF was drawn in. The collection of CSF was allowed to occur slowly. Once the desired volume was collected, the tubing’s three-way valve was closed to stop the flow of CSF. The capillary was gently removed using the micromanipulator. A collection tube (1.5 mL microcentrifuge tube with 1 µl protease inhibitor) was positioned beneath the capillary’s tip. Upon opening the tubing, the syringe plunger was slowly pressed to allow the CSF to flow into the collection tube. After the collection process, the CSF was pooled together (n=8/group) and briefly centrifuged. The CSF samples were analyzed by Western blot analysis with an anti-ASRA antibody (Proteintech, cat#30246-1-AP). Purified rASRA protein was used as a standard.

### Measurement of circulating glucose, insulin and leptin

Glucose level was measured using a glucose meter. Commercial ELISA kits were used to measure insulin (CrystalChem, cat#90080) and leptin (CrystalChem, cat#90030) levels, which were calculated with four-parameter logistic curve fit. In some experiments, due to large cohorts, leptin level was measured in a randomly picked subset of serum samples.

### Adipocyte culture and differentiation

The immortalized brown preadipocyte cell line was previously generated^69^. On day −2 of differentiation, brown preadipocytes at 70% confluence were cultured in Dulbecco’s Modified Eagle’s Medium (DMEM, cat#11965-092, Gibco) supplemented with 10% fetal bovine serum (FBS, cat#S11550, Atlanta biologicals), 20 nM insulin (Sigma, cat#I6634), 1 nM 3,3′,5-triiodo-l-thyronine (Sigma, cat#T0281), 50 units/ml penicillin, and 50 mg/ml streptomycin (differentiation medium). To induce adipocyte differentiation on day 0, the cells were cultured in differentiation medium supplemented with 0.125 mM indomethacin (Alfa Aesar, cat#A19910-06), 0.5 μM dexamethasone (Sigma, cat#d4902), and 0.5 mM isobutylmethylxanthine (Sigma, cat#I7018) for 48 hours. After the induction period, the cells were returned to the differentiation medium. The primary iWAT preadipocyte culture and differentiation were described previously^18, 19^. Preadipocytes were isolated from two-week-old mice and cultured until confluence. Differentiation was initiated by culturing the confluent cells in DMEM/F12 medium (Gibco, cat#11320-033) containing 10% FBS, 850 nM insulin, 1 nM triiodothyronine, 0.5 mM isobutylmethylxanthine, 0.5 μM dexamethasone, and 0.125 mM indomethacin. After 2 days, the cells were maintained in DMEM/F12 medium supplemented with 10% FBS, 850 nM insulin, and 1 nM triiodothyronine, with the medium being changed every 2 days. On day 6, both brown and primary iWAT adipocytes were fully differentiated, and cells with at least 95% differentiation efficiency were used in experiments.

### ASRA secretion

Differentiated adipocytes were cultured in serum-free DMEM medium for 6 hr. Cell extracts and conditioned medium were collected and subjected western blot analysis using an antibody against ASRA (Santa Cruz Biotechnology, discontinued, cat#sc-163566). The specificity of the antibody was validated. To examine ASRA secretion *ex vivo*, BAT tissue was cut into small pieces and incubated with serum-free DMEM medium. At indicated time points, medium was collected and replenished with fresh serum-free DMEM medium. To examine ASRA secretion in HEK293 cells, cells were transfected with ASRA-GFP, FGF1-GFP, or GFP expression vector alone. Two days after transfection, cells were cultured in either serum-free DMEM medium containing 4.5 g/L glucose or serum-free low glucose (1 g/L) DMEM medium for 6 hr. Medium was collected and GFP signal intensity was measured by a Synergy H4 Hybrid microplate reader (BioTek, Winooski, VT) using Gen5 software. Background signal (culture medium from cells without plasmid transfection), which was similar to that of GFP medium, was subtracted.

### ASRA subcellular localization

To investigate endogenous ASRA localization, mature adipocytes were cultured in serum-free DMEM medium containing either 4.5 g/L or 1 g/L glucose for 6 hr. The cells were then fixed with 3.7% formalin for 15 minutes at room temperature. After fixation, permeabilization was achieved by exposing the cells to ice-cold 100% methanol at −20°C for 10 minutes, followed by washing twice with PBS. Subsequently, the cells were incubated in blocking buffer (PBS containing 5% normal goat serum and 0.3% Triton X-100) at room temperature for 60 minutes. The cells were then subjected to overnight incubation at 4°C with a 1:100 dilution of ASRA antibody (Sigma, cat#HPA049367) and a 1:500 dilution of LC3 antibody (MBL, cat#M152-3) in the blocking buffer. The ASRA antibody was validated. After incubation, cells underwent gentle PBS washing twice and were incubated with Alexa Fluor goat anti-rabbit 488-conjugated secondary antibody (ThermoFisher, cat#A-11008) and Alexa Fluor goat anti-mouse 594-conjugated secondary antibody (ThermoFisher, cat#A-11032) for 2 hours at room temperature. Finally, cells were exposed to DAPI at room temperature for 30 minutes. The images were obtained utilizing an inverted Nikon Eclipse Ti2 confocal microscope (Nikon Instruments/Nikon Corp) and were subjected to uniform processing settings.

To investigate ASRA-GFP localization in mature adipocytes and COS7 cells, cells were co-transfected with ASRA-GFP plasmids along with mCherry-ER, RFP-Golgi, or RFP-LC3 plasmids. Following a 48-hour post-transfection period, the cells were fixed using 3.7% formalin for 15 minutes at room temperature. Subsequently, DAPI (Sigma, cat#D9542) was applied and incubated at room temperature for 30 minutes to stain the nuclei. The images were obtained and processed as described above.

### Gene expression

Total RNA was extracted from mature adipocytes or tissues using TRIzol reagent (Invitrogen, cat# 15596-018) following the manufacturer’s instructions. An equal amount of RNA was used for reverse transcription. Quantitative real-time PCR (qRT-PCR) was performed using SYBR green fluorescent dye (Bio-Rad, cat#1725272) on an ABI7300 PCR instrument. The ribosomal 36B4 (U36) gene was used as an internal control. The relative mRNA expression levels were calculated using the ΔΔ^-Ct^ method. Primer sequences will be provided upon request.

### Bioinformatics analysis of public gene expression datasets

We analyzed microarray data (GSE7623)^20^ for fasting-induced genes. We downloaded and analyzed the RNA-seq data from GSE86338 dataset^70^ using DESeq2, then plotted WAT ASAR mRNA expression during room temperature and chronic cold exposure. We downloaded and analyzed the RNA-seq data from GSE88818 dataset^71^ using DESeq2, and then plotted liver ASRA mRNA expression under normal chow diet and HFD. The linear regression between ASAR and BMI was analyzed on microarray data (GSE70353)^23^ from subcutaneous adipose tissue samples of a cohort of 770 men using GraphPad Prism 7.

### Statistical Analysis

The sample size for this study was determined based on prior experience and existing literature in the field. The investigators conducting the mouse experiments were not blinded to the genotypes. The number of biological samples (n) was provided for each figure panel. Unless otherwise specified, the data were presented as mean ± standard error of the mean (s.e.m.), and individual data points were plotted. Statistical analyses were performed using GraphPad Prism 8.0 software. Analysis between 2 groups was performed using Student’s t-test. Time course analysis was done by two-way ANOVA followed by a post-hoc test using Bonferroni’s method for individual time points. Statistical significance was considered when P < 0.05.

## Reporting summary

Further information on research design is available in the Nature Portfolio Reporting Summary linked to this article.

## Data availability

All data in the article and supplementary information are available. Any additional information required for the data reported in this paper is available upon reasonable request.

