## Supplemental Figures for "A brown fat-enriched adipokine, ASRA, is a leptin receptor antagonist that stimulates appetite"

### Extended Data Figure 1

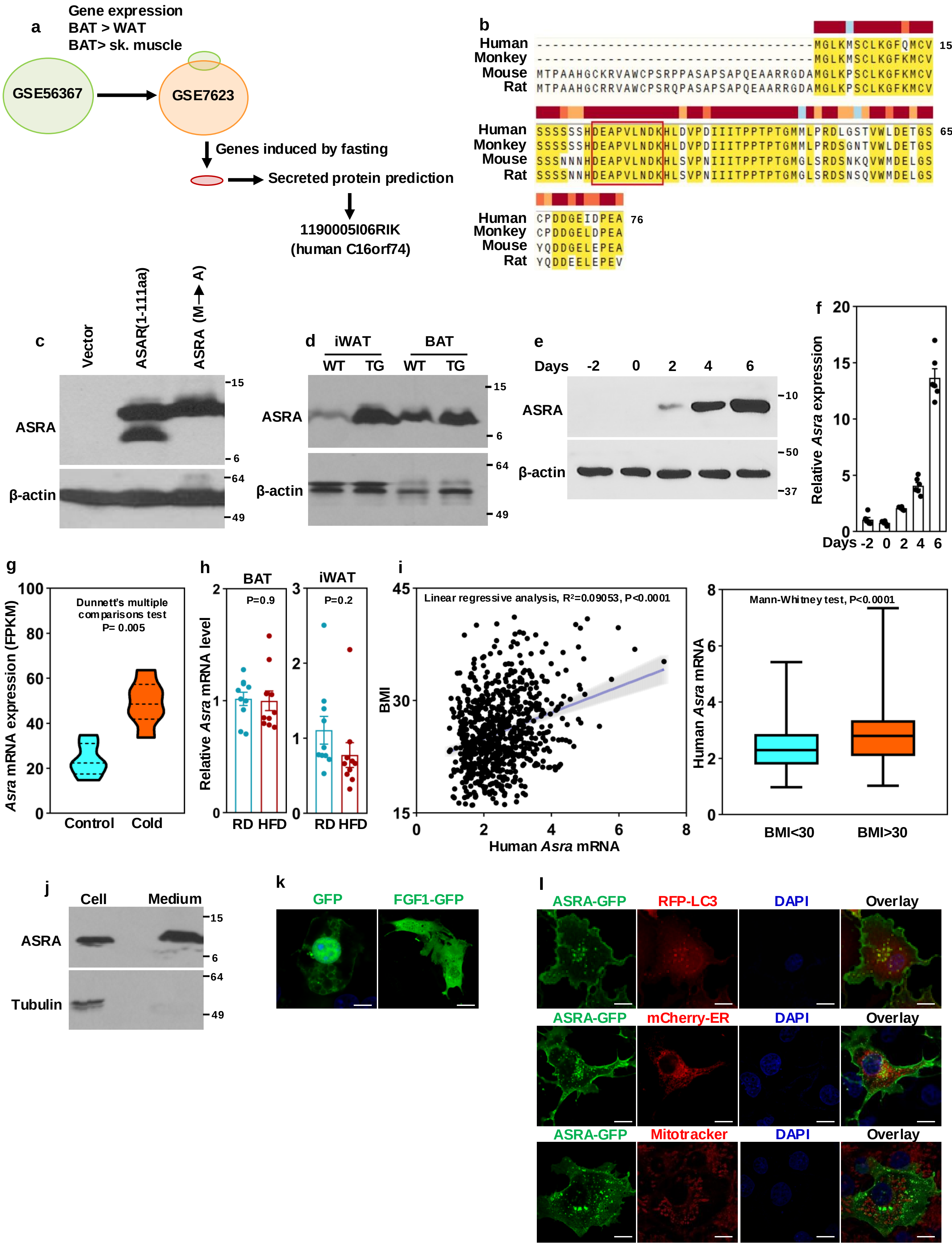

**Extended Data Figure 1**

**a,** Strategies to identify BAT-secreted ASRA.

**b,** Human, monkey, mouse, and rat ASRA protein sequences.

**c,** ASRA protein in HEK293 cells transfected with indicated plasmids.

**d,** ASRA protein expression in iWAT and BAT of wild type (WT) and aP2-*Asra* transgenic mice (TG).

**e, f,** ASRA protein (**e**) and mRNA (**f**) expression during adipocyte differentiation.

**g,** *Asra* mRNA expression in WAT of mice at room temperature (n=4) and 4$^{o}$C (n=6) for seven days. The RNA-seq data were downloaded from GSE86338 dataset.

**h,** *Asra* mRNA expression in BAT and iWAT fed with regular diet (RD) or high-fat diet (HFD). n=10/group.

**i,** Analysis of *Asra* mRNA expression in subcutaneous WAT of 770 men in relation to BMI. The microarray data were downloaded from GSE70353 dataset.

**j,** ASRA secretion in mature adipocytes.

**k,** FGF1-GFP localization in mature adipocytes. Bar=50 μm.

**l,** Co-localization of ASRA-GFP with RFP-LC3 in COS7 cells. Bar=100 μm.

### Extended Data Figure 2

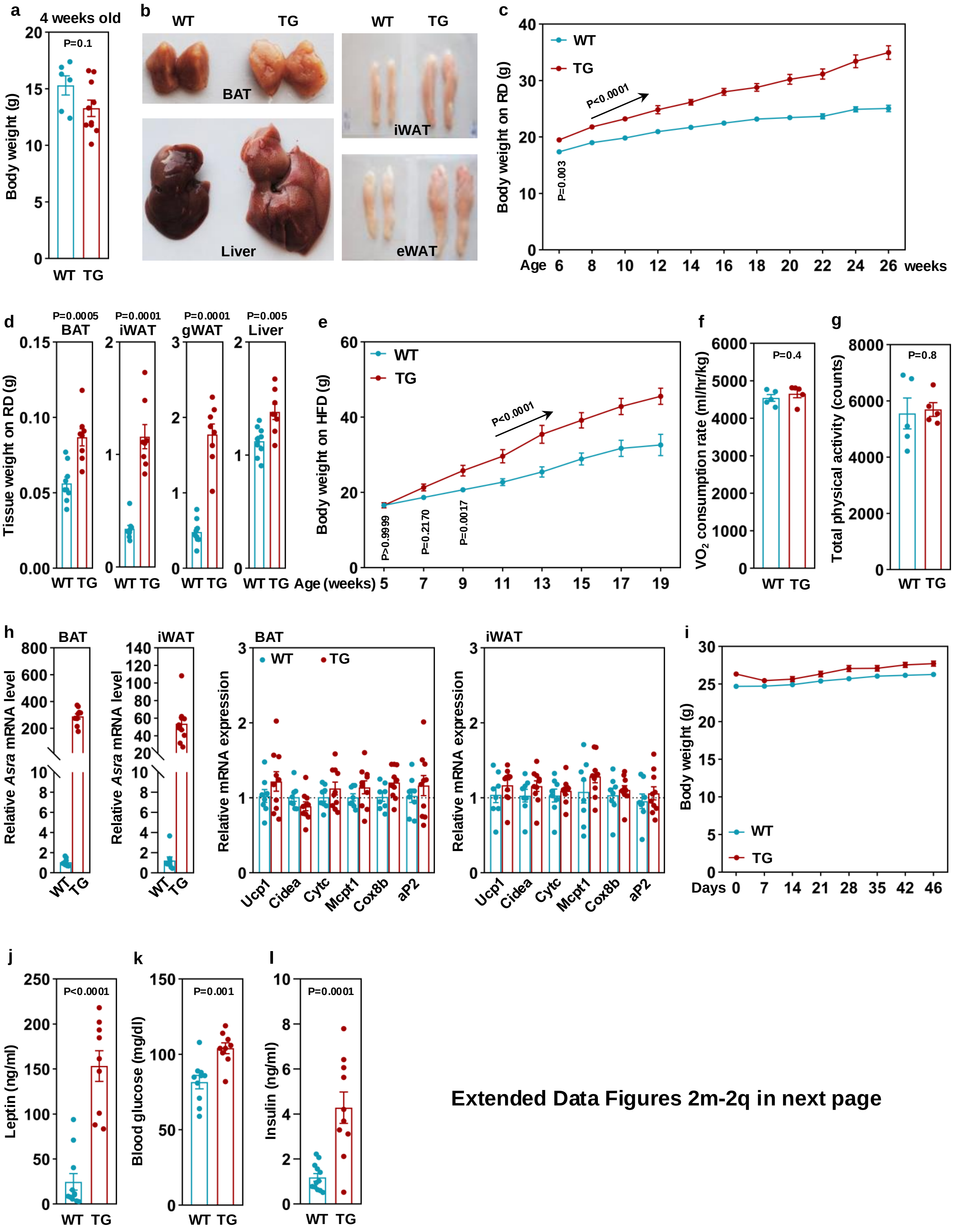


**Extended Data Figures 2m-2q in next page**

### Extended Data Figure 2 (continued)

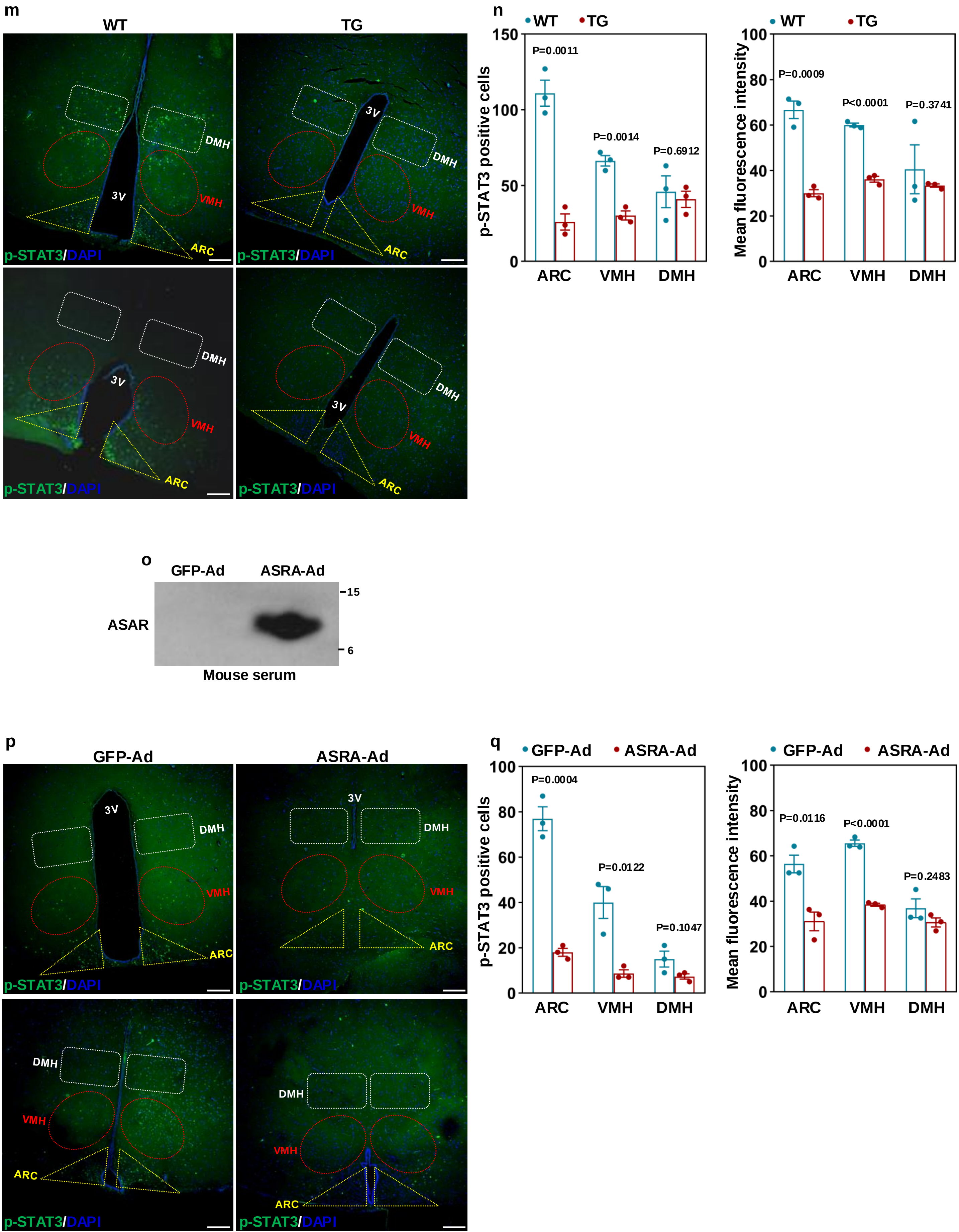

**Extended Data Figure 2**

**a,** Body weight of male aP2-*Asra* TG (n=10) and littermate controls (n=6) at four-weeks-old.

**b,** Representative tissue images of mice on a regular diet.

**c,** Body weight of female aP2-*Asra* TG mice (n=12) and littermate controls (n=9) on a regular diet.

**d,** Tissue weight of female aP2-*Asra* TG mice (n=8) and littermate controls (n=9) on a regular diet.

**e,** Body weight of female aP2-*Asra* TG mice (n=6) and littermate controls (n=7) on a HFD.

**f, g,** Oxygen consumption (**f**) and physical activity (**g**) assessed by metabolic cages studies. n=5/group.

**h,** Gene expression in BAT and iWAT of 2-month-old male aP2-*Asra* TG mice (n=10) and littermate controls (n=8).

**i,** Body weight of male aP2-*Asra* TG mice (n=10) and littermate controls (n=6) during a pair-feeding experiment.

**j, k, l,** Circulating leptin (**j**), glucose (**k**) and insulin (**l**) levels in aP2-*Asra* TG mice and littermate controls at 10-month-old. n=9-12/group.

**m,** Phospho-STAT3 immunostaining in the ACR, VMH and DMH of hypothalamus in aP2-*Asra* TG mice and littermate controls that were fasted for 5 hr followed by leptin injection. Bar=200 μm.

**n,** Quantification of phospho-STAT3-positive cells and mean fluorescence intensity in hypothalamic ARC, VMH and DMH regions (n=3).

**o,** Detection of ASRA in serum of mice injected with ASRA adenoviruses.

**p,** Phospho-STAT3 immunostaining in the ACR, VMH and DMH of hypothalamus in adenovirus-infected mice that were fasted for 5 hr followed by leptin injection. Bar=200 μm.

**q,** Quantification of pSTAT3-positive cells and mean fluorescence intensity in hypothalamic ARC, VMH and DMH regions (n=3).

#### Extended Data Figure 3

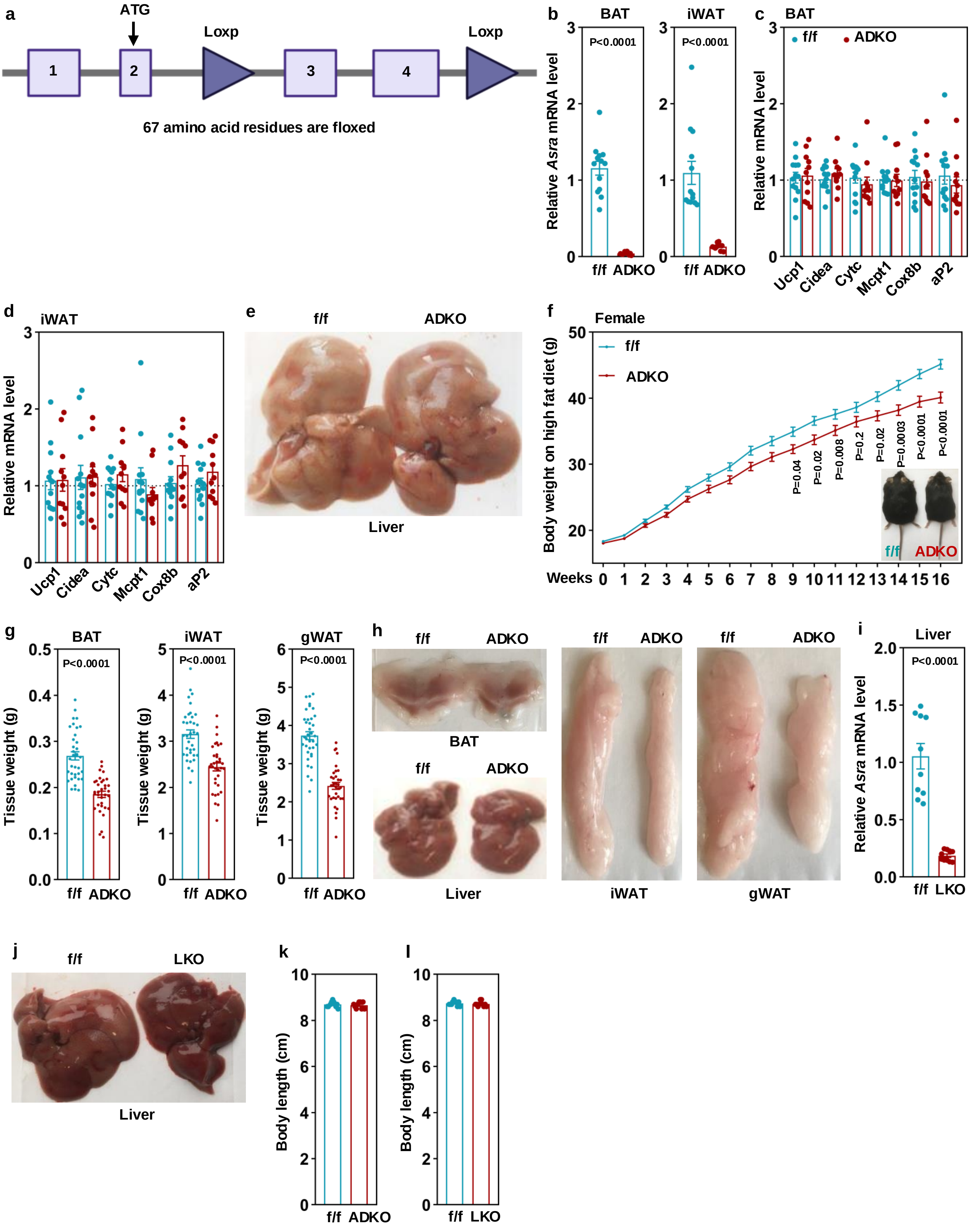

**Extended Data Figure 3**

**a**, Exon 3 and 4 of *Asra* were floxed by two LoxP sites.

**b, c, d,** Gene expression in BAT and iWAT from 2-month-old male *Asra* ADKO mice (n=11) and littermate controls (n=13).

**e,** Representative liver images of *Asra* ADKO male mice and littermate controls on a HFD.

**f,** Body weights of female *Asra* ADKO mice (n=34) and littermate controls (n=36) on a HFD.

**g,** Tissue weights of mice in (**f**). gWAT, gonadal WAT. n=34-36/group.

**h,** Representative tissue images of mice in (**f**).

**i,** *Asra* expression in liver of 2-month-old male *Asra* LKO mice (n=10) and littermate controls (n=10).

**j,** Representative liver images of *Asra* LKO male mice and littermate controls on a HFD.

**k,** Body length of male *Asra* ADKO (n=9) and littermate controls (n=12) at three-month-old.

**l**, Body length of male *Asra* LKO (n=10) and littermate controls (n=10) at three-month-old.

### Extended Data Figure 4

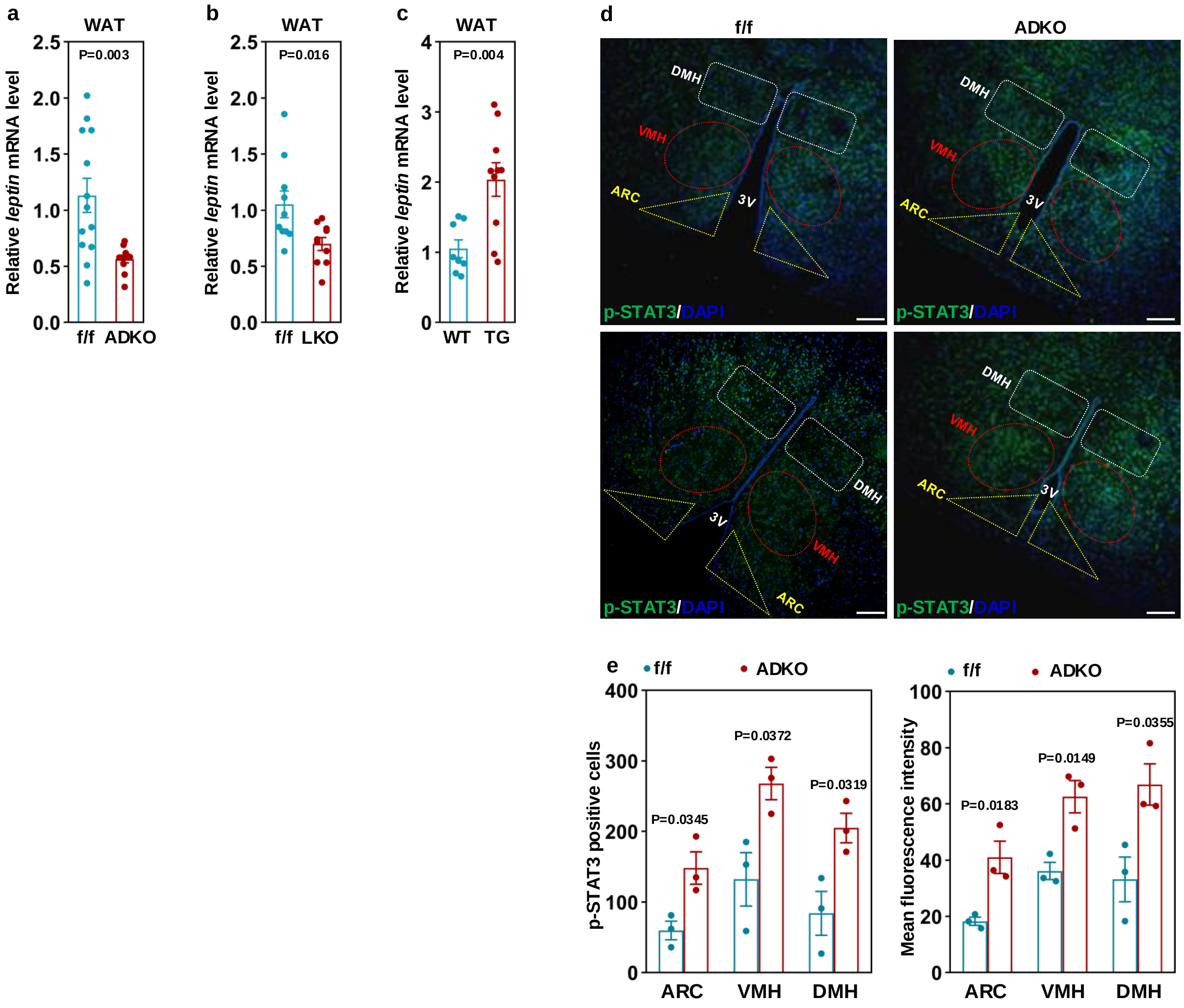


Extended Data Figures 4f-4i in next page

### Extended Data Figure 4 (continued)

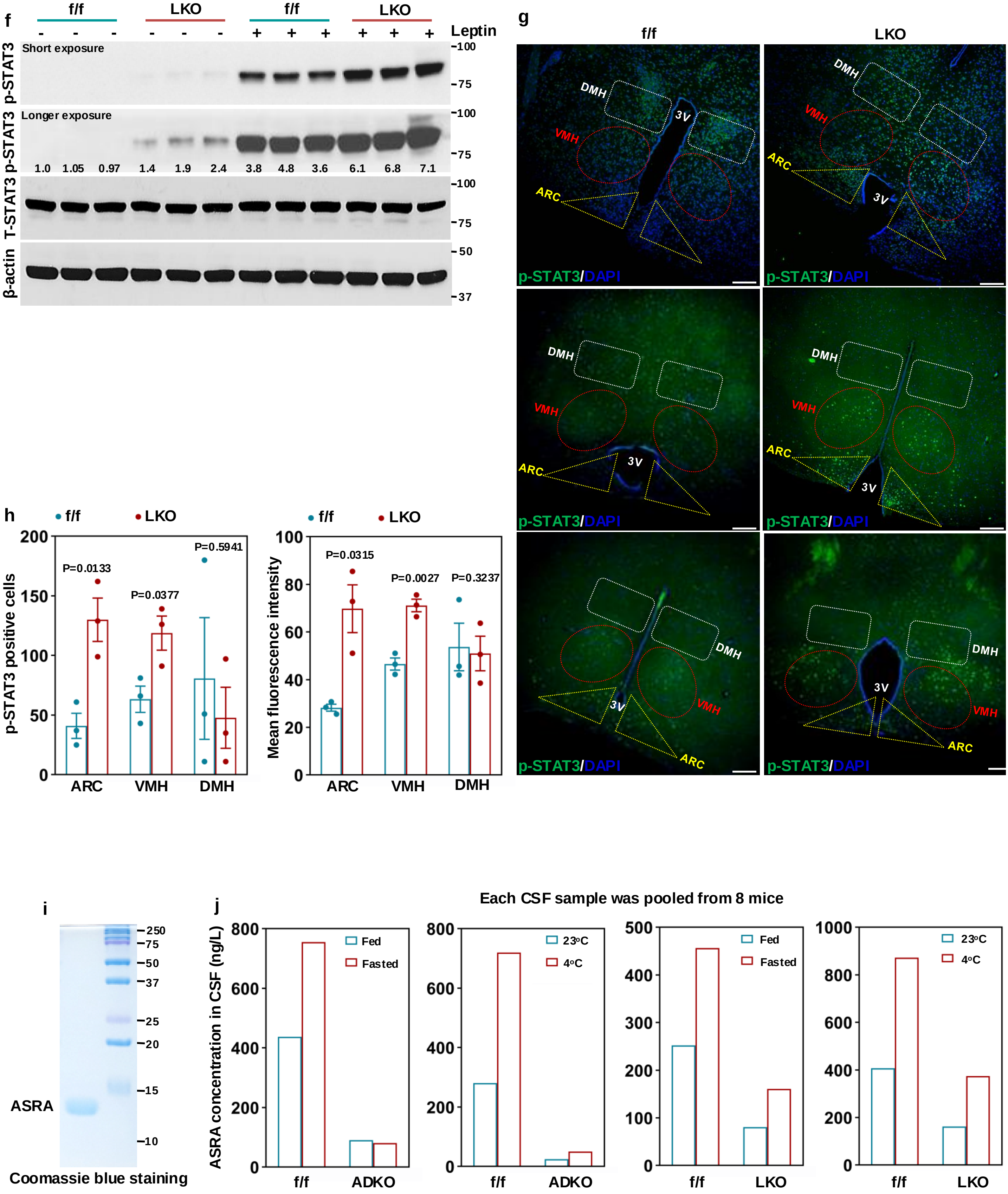

**Extended Data Figure 4**

**a, b,** *Leptin* mRNA expression in WAT of 3-month-old male *Asra* ADKO (n=11) and littermate controls (n=13) (**a**), and 3-month-old male LKO (n=10) and littermate controls (n=10) (**b**).

**c,** *Leptin* mRNA expression in WAT of 3-month-old male aP2-*Asra* TG (n=10) and littermate controls (n=8).

**d,** Phospho-STAT3 immunostaining in the ARC, VMH and DMH of hypothalamus in *Asra* ADKO mice and littermate controls that were fasted for 5 hr followed by leptin injection. Bar=200 μm.

**e,** Quantification of phospho-STAT3-positive cells and mean fluorescence intensity in hypothalamic ARC, VMH and DMH regions (n=3).

**f,** Phosphorylation of STAT3 in hypothalamus of LKO mice and littermate controls that were pre-fasted for 3 hr.

**g,** Phospho-STAT3 immunostaining in the ARC, VMH and DMH of hypothalamus in *Asra* LKO mice and littermate controls that were fasted for 5 hr followed by leptin injection. Bar=200 μm.

**h,** Quantification of phospho-STAT3-positive cells and mean fluorescence intensity in hypothalamic ARC, VMH and DMH regions (n=3).

**i,** Coomassie blue staining of rASRA protein purified from Expi293 cells.

**j,** ASRA concentrations in CSF were estimated from Western blots. Each CSF sample was pooled from 8 mice.

### Extended Figure 5

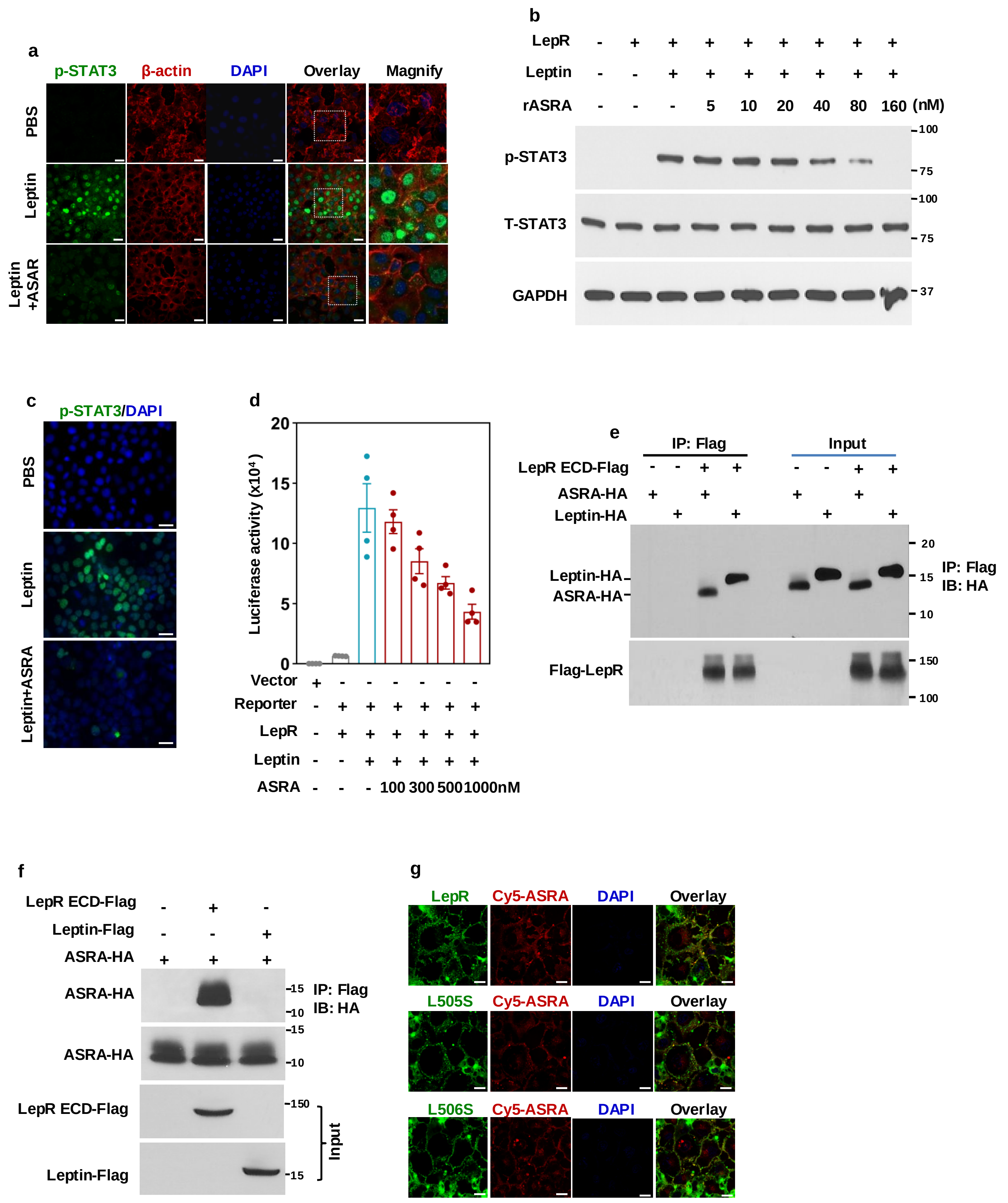

**Extended Data Figure 5**

**a,** Phospho-STAT3 immunostaining in COS7 cells transfected with the human leptin receptor. Cells were treated with rASRA (300 nM) for 30 min and leptin (100 nM) was added for another 30 min. Bar=100 μm.

**b,** Western blot analysis of phospho-STAT3 of HEK293 cells treated with rASRA and leptin as described in (**a**).

**c,** Phospho-STAT3 immunostaining in COS7 cells transfected with the human leptin receptor. Cells were treated with rASRA (300 nM) purified from bacteria for 30 min and leptin (100 nM) was added for another 30 min. Bar=100 μm**.**

**d,** HEK293 cells were transfected with indicated plasmids, treated with leptin (100 nM) and rASRA purified from bacteria at different concentrations for 24h, and the luciferase activity was measured. n=4.

**e, f,** HEK293 cells were transfected with indicated plasmids and the conditioned medium were immunoblotted with an HA antibody after immunoprecipitation with a Flag antibody.

**g,** COS7 cells transfected with human leptin receptor mutants were incubated with Cy5-labeled rASRA protein (300 nM) for 30 min. Bar=100 μm.

#### Extended Figure 6

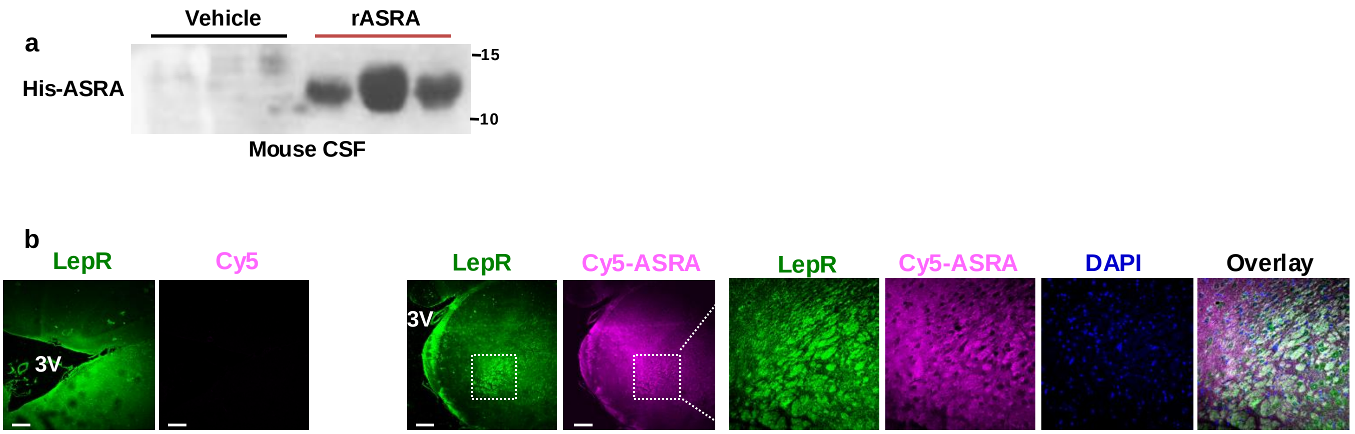

**Extended Data Figure 6**

**a,** Three-month-old male mice were ip injected with His-tagged rASRA protein (5.4 nmole/mouse). The presence of rASRA in CSF was detected with a His-tag antibody.

**b,** Three-month-old male mice were tail-vein injected with Cy5 dye (24 nmole/mouse) or Cy5-ASRA (5.4 nmole/mouse). Two-hour post injection, the binding of Cy5-ASRA was examined and leptin receptor immunostaining was examined in hypothalamic slices.
